# Massive computational acceleration by using neural networks to emulate mechanism-based biological models

**DOI:** 10.1101/559559

**Authors:** Shangying Wang, Kai Fan, Nan Luo, Yangxiaolu Cao, Feilun Wu, Carolyn Zhang, Katherine A. Heller, Lingchong You

**Affiliations:** Department of Biomedical Engineering, Duke University, Durham, NC 27708; Department of Statistical Science, Duke University, Durham, NC 27708; Center for Genomic and Computational Biology, Duke University, Durham, NC, 27708; Molecular Genetics and Microbiology, Duke University, Durham, NC, 27708

## Abstract

Mechanism-based mathematical models are the foundation for diverse applications. It is often critical to explore the massive parametric space for each model. However, for many applications, such as agent-based models, partial differential equations, and stochastic differential equations, this practice can impose a prohibitive computational demand. To overcome this limitation, we present a fundamentally new framework to improve computational efficiency by orders of magnitude. The key concept is to train an artificial neural network using a limited number of simulations generated by a mechanistic model. This number is small enough such that the simulations can be completed in a short time frame but large enough to enable reliable training of the neural network. The trained neural network can then be used to explore the system dynamics of a much larger parametric space. We demonstrate this notion by training neural networks to predict self-organized pattern formation and stochastic gene expression. With this framework, we can predict not only the 1-D distribution in space (for partial differential equation models) and probability density function (for stochastic differential equation models) of variables of interest with high accuracy, but also novel system dynamics not present in the training sets. We further demonstrate that using an ensemble of neural networks enables the self-contained evaluation of the quality of each prediction. Our work can potentially be a platform for faster parametric space screening of biological models with user defined objectives.

## Introduction

Mathematical modeling has become an indispensable tool in analyzing the dynamics of biological systems at diverse length- and time-scales^1–6^. In each case, a model is typically formulated to account for the biological processes underlying the system dynamics of interest. When analyzing a gene circuit, the corresponding model often entails description of the gene expression; for a metabolic pathway, the corresponding model may describe the constituent enzymatic reactions; for an ecosystem, the corresponding model would describe growth, death, and movement of individual populations, which could in turn be influenced by other populations. We call these models mechanism-based models.

Mechanism-based models are useful for testing our understanding of the systems of interest^7–14^. For instance, modeling has been used to examine of the network motifs or the parameter sets able to generate oscillations^15,16^ or spatial patterns^17^, or the noise characteristics of signaling networks^18–21^. They may also serve as the foundation for practical applications, such as designing treatments of diseases^22–24^ and interpreting the pharmacokinetics of drugs^25–27^. Many mechanism-based models cannot be solved analytically and have to be analyzed by numerical methods. This situation is particularly true for models dealing with spatial or stochastic dynamics. While numerical simulations are typically more efficient than experiments, they can still become computationally prohibitive for certain biological questions. For example, consider a model with 10 parameters. To examine 6 values per parameter, there will be 6^10^ parameter combinations. If each simulation takes 5 minutes, which is typical for a PDE model, the screening would require 575 years to finish. Many biological systems are much more complex. For each system, both the size of the parametric space and the time required to do each simulation would increase combinatorically with the system complexity. Thus, standard numerical simulations using mechanism-based models can face a prohibitive barrier for large-scale exploration of system behaviors.

Thanks to its ability to make predictions without a full mapping of the mechanistic details, deep learning has been used to emulate time-consuming model simulations^28–31^. To date, however, the predicted outputs are restricted in categorical labels or a set of discrete values. By contrast, deep learning has not been used to predict outputs consisting of continuous sequences of data (e.g., time series, spatial distributions, and probability density functions). We overcome this limitation by adopting a special type of deep learning network, the LSTM (Long-Short-Term Memory) network. Our approach leads to ~30,000-fold acceleration in computation with high prediction accuracy (See network performance in Methods for estimation details). We further develop a voting strategy, where multiple neural networks are trained in parallel, to assess and improve the reliability of predictions.

## Results

### The conceptual framework

When numerically solving a mechanism-based dynamic model consisting of differential equations, the vast majority of the time is spent in the generation of time courses. For many biological questions, however, the main objective is to map the input parameters to specific outcomes, such as the ability to generate oscillations or spatial patterns^32–37^. For such applications, the time-consuming generation of time courses is a “necessary evil”.

The key to the use of the deep learning is to establish this mapping through training to bypass the generation of time courses, leading to a massive acceleration in predictions (Figure 1). To do the learning, we use a small proportion of data generated by the mechanism-based model to train a neural network. The data generated by the mechanistic model need to be sufficiently large to ensure reliable training but small enough such that the data generation is computational feasible.

**Figure 1.**
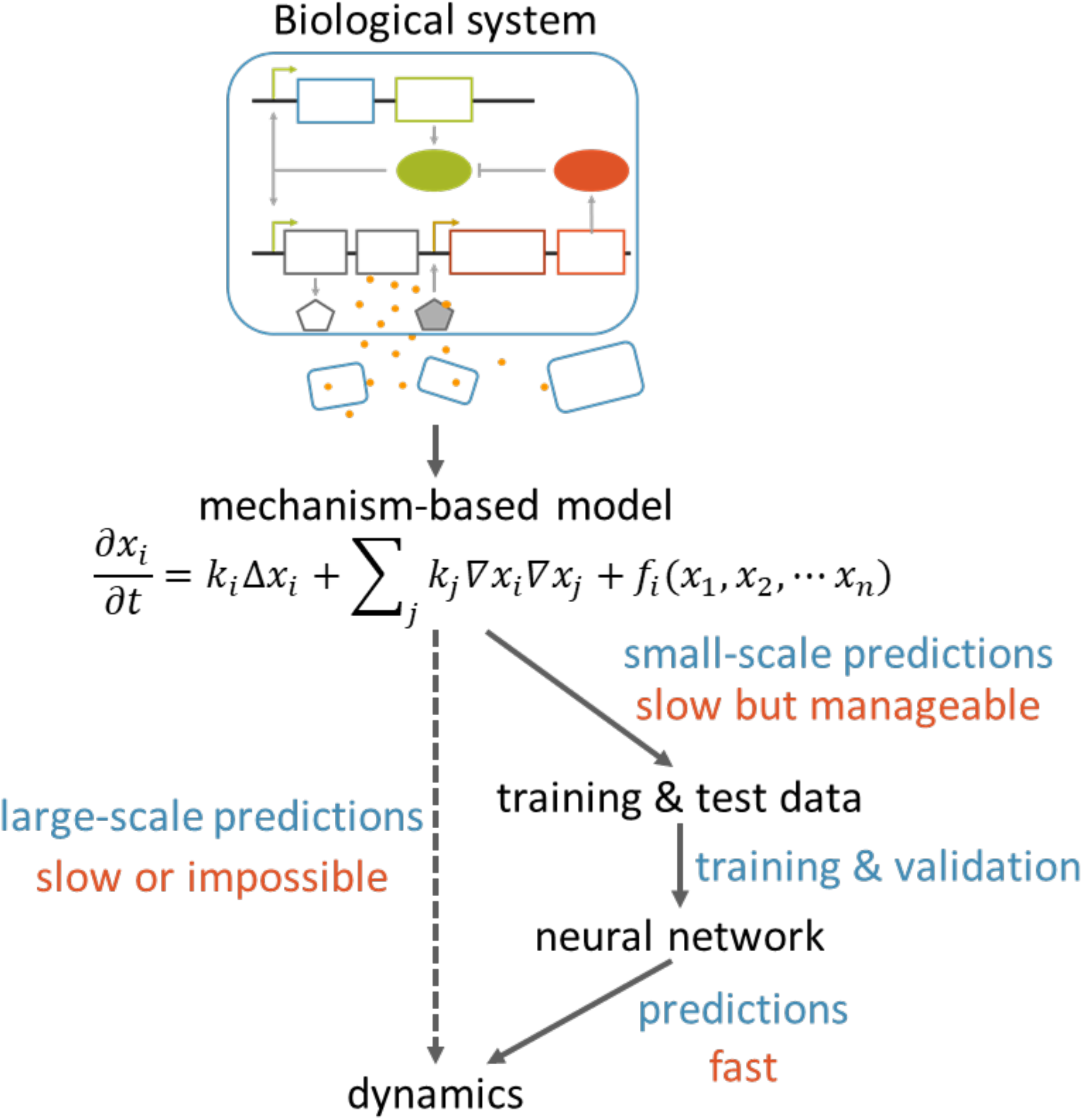
Using an artificial neural network to emulate a mechanism-based model. Here a hypothetic biological network and the corresponding mechanistic model are shown. The mechanistic model is used to generate a training data set, which is used to train a neural network. Depending on the specific mechanistic model, the trained neural network can be orders of magnitude faster, enabling exploration of a much larger parametric space of the system.

As a proof of principle, we first apply our approach to a well-defined model developed by Cao et al.^32^ This PDE model describes pattern formation in *Escherichia coli* programed by a synthetic gene circuit (<u>Methods</u> and Figure S1A), accounting for cell growth and movement, intercellular signaling and circuit dynamics as well as transportation (Equation 1). This model was previously used to capture the generation of characteristic core-ring patterns and to examine the scaling property of these patterns. Numerical simulations were used to explore the parametric space to seek parameter combinations able to generate scale-invariant patterns. Several months were needed to search through 18,231 parameter sets. Yet, these parameter sets only represent an extremely tiny fraction of the parametric space that the system can occupy. Thus, it is likely that these numerical simulations have not revealed the full capability of the system in terms of pattern formation. For example, it is unclear whether the system can generate multiple rings and how this can be achieved.

For this system, each input is a set of parameters (e.g., cell growth rate, cell motility, and kinetic parameters associated with gene expression); the output is the spatial distribution of a molecule. The mapping between the two is particularly suited for the use of an LSTM network. The LSTM network, a type of recurrent neural network (RNN), was proposed in 1997 to process outputs consisting of a continuous series of data^38^. It has demonstrated great potential in natural language processing and speech recognition as well as in other sequence-prediction applications^39^.

The outputs of the model can vary drastically in the absolute scale. To improve the learning process, we break each output profile into two components: the peak value of each profile and the profile normalized with respect to the peak value. Our deep neural network consists of an input layer with inputs to be the parameters of mechanism-based model, connected to a fully connected layer, and the output layer consists of two types of outputs, one for predicting peak value of the profile, directly connected to the fully connected layer, the other for predicting the normalized profile, connected to LSTM cell arrays, which was fed by the output from fully connected layer. The detailed structure of the neural network is described in <u>Methods</u> and Figure S2.

### The neural network predicts spatial distributions with high accuracy

To train the neural network, we first used our PDE model to generate 10^5^ simulation results from random combinations of parameter values. Generating these data sets was manageable: it took 2 months on a cluster consisting of 400 nodes. We split the data sets into three groups: 80% for training, 10% for validation and 10% for testing. We used the mean-squared error (MSE) to evaluate the difference between the data generated by PDE simulation and those generated by the neural network. Each output distribution was dissected into two components for prediction: the peak value, and the shape of the distribution (i.e. the distribution after being normalized with respect to the peak value).

The trained neural network is highly accurate. The correlation between predicted values and PDE simulation results exhibits high *R^2^* values: 0.988 for peak value predictions and 0.997 for shape predictions (Figure 2A). For most parameter sets, distributions generated by the neural network align nearly perfectly with those generated by numerical simulations (Figure 2B, S3, and S4). In general, the more complex the output distribution, the less accurate the prediction (Table S4). This trend likely results from the uneven representation of different types of patterns in the training data sets: the majority have no ring (42897 out of 10^5^) or have only 1 ring (55594 out of 10^5^); patterns with multiple rings are rare (1509 out of 10^5^ for 2 rings or more) (Figure S5).

**Figure 2.**
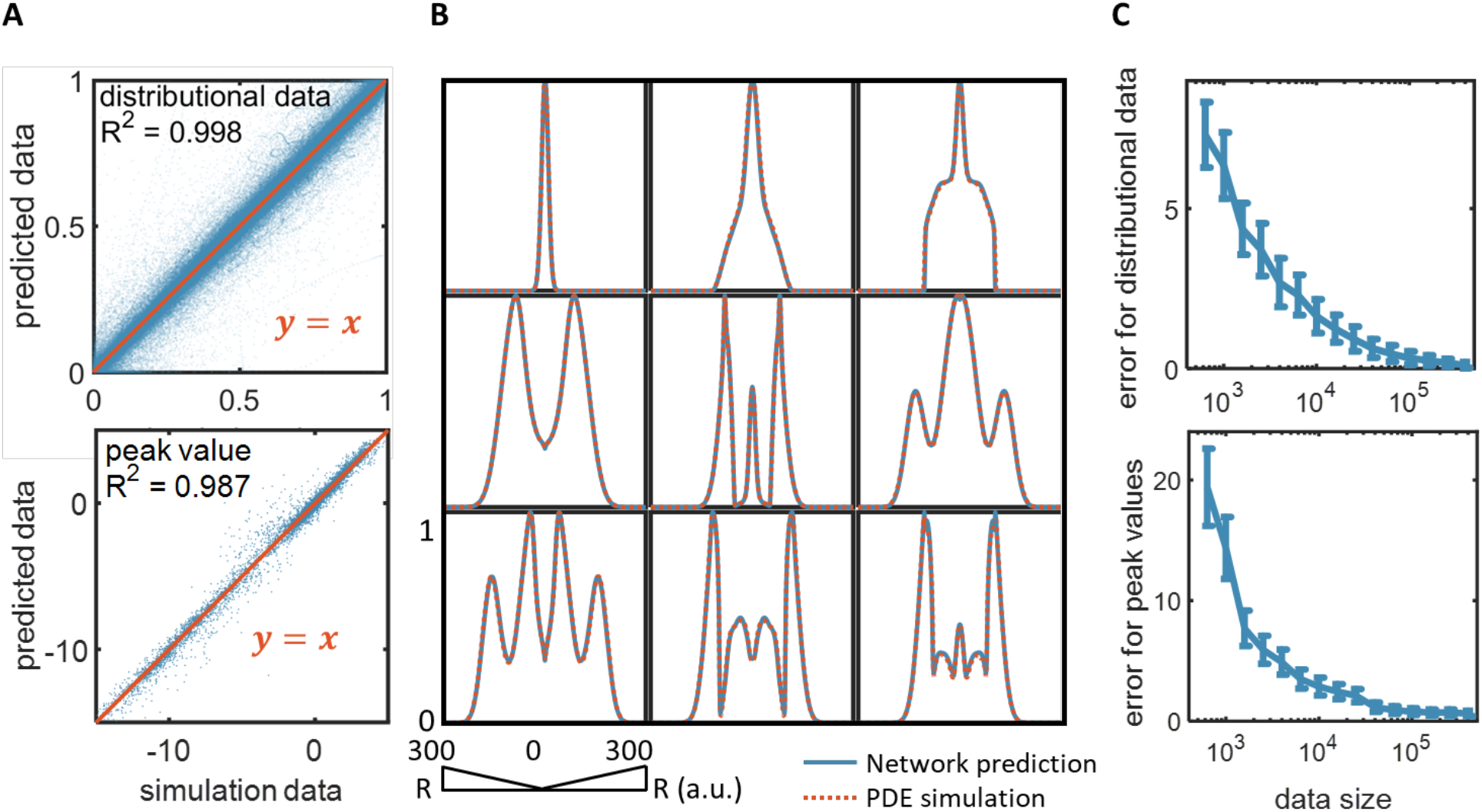
Neural network training and performance. We generated 10^5^ simulated spatial distributions using our PDE model and split the data into three groups: 80% for training, 10% for validation and 10% for test. We used MSEs to evaluate the differences between data generated by the mechanism-based model and data generated by the neural network. **A. Accuracy of the trained neural network.** The top panel shows the predicted distributions by the neural network plotted against the distributions generated by numerical simulations. The bottom panel shows the peak values predicted by the neural network plotted against the peak values generated by numerical simulations. Perfect alignment corresponds to the *y* = *x* line. The test sample size is s (=10,000). Each spatial distribution consists of 501 discrete points; thus, the top panel consists of 5,010,000 points. **B. Representative distributions predicted by neural network.** Each blue line represents a predicted distribution using the trained neural network; the corresponding red dashed line represents that generated by a numerical simulation. Additional examples are shown in Figures S3 and S4. **C. Identifying the appropriate data size for reliable training.** The top panel shows the MSE between distributions generated by the neural network and the distributions generated by numerical simulations as a function of an increasing training data size. The bottom panel shows the MSE of peak-value predictions as a function of an increasing training data size. Unless noted otherwise, we use a training data size of 10^5^, where the MSEs are sufficiently small. The MSEs are calculated based on predictions of a test dataset, which contains 20,000 samples.

To identify the minimum size of dataset needed for accurately making predictions, we trained deep LSTM network on different training dataset sizes. The MSEs are calculated based on predictions of a fixed test dataset, which contains 20,000 samples. Figure 2C demonstrates how the MSEs of distributional data and peak values decrease with the increase of training data size. Since the x-axis is log-scaled, when the dataset size is beyond 10^4^, the rate of error reduction becomes asymptotically smaller. When the data size is 10^5^, the MSE is sufficiently small. This training dataset size is manageable and results in sufficient accuracy for our analysis. Based on error tolerance and numerical data generation efficiency, one can choose the desired dataset size for training the neural network. With an ensemble of deep neural networks, which will be described in the next section, the errors can be further reduced without increasing the dataset size.

### The neural network predicts novel patterns

We use trained deep LSTM network to screen through the parametric space. It takes around 12 days to screen through 10^8^ combination of parametric sets, which would need thousands of years if generated with PDE simulations. We find more than 1000 three-ring pattern distributions, including novel patterns not present in the training sets (Figure 3A). These are genuinely novel 3-ring patterns found in this screening process, which are not in the training dataset. We further validate them by numerical simulations using the PDE model and find rare exceptions (Figure 3B, Table S3). Their existence indicates that the learning by the neural network is not limited to passive recollection of what the network has been trained with. Instead, the training has enabled the neural network to establish the genuine mapping between the input parameters and the system outputs, in a manner that is highly non-intuitive.

**Figure 3.**
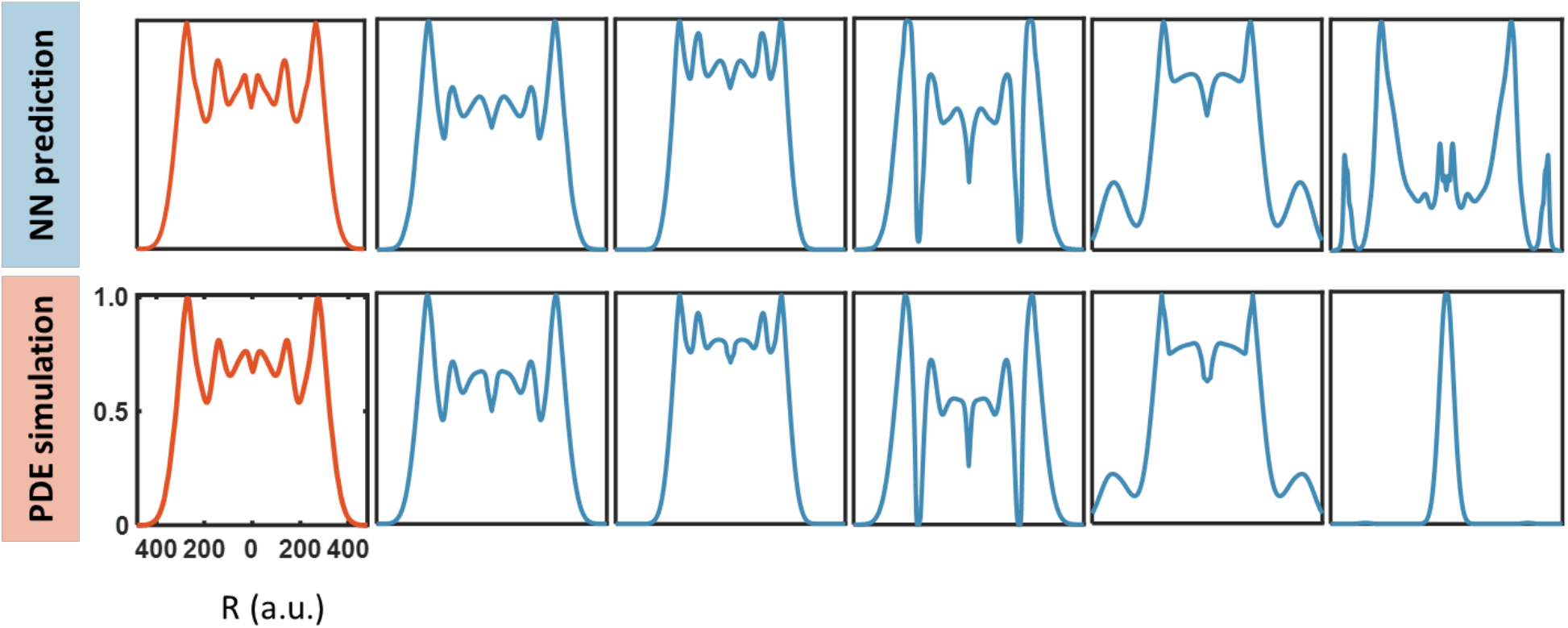
The trained neural network predicts novel patterns. We used the neural network to screen 10^8^ parameter combinations to search for 3-ring patterns. We then use the mechanism-based model to test accuracy of predicted patterns (We test 1284 3-ring patterns and the mean value of the MSEs between NN predicted distributions and PDE simulations is 0.079 and the standard deviation is 0.008). The distributions shown in red are from training data set. The other distributions are from the test data sets (top) and the corresponding results generated by the mechanism-based model for validation (bottom). In four examples, the neural network predictions are validated. In one, the neural prediction is incorrect.

### Voting between neural networks enables estimation of prediction accuracy

Despite the extremely high accuracy in the predictive power of the trained neural network, it is never 100% correct. This apparent deficiency is the general property of neural networks. The lack of perfection in prediction raises a fundamental question: when dealing with a particular prediction, how do we know it is sufficiently reliable? Even if it were feasible, validating every prediction by simulation, as done in the last section (Figure 3), *would defeat the purpose of using deep learning*. Therefore, it is critical to develop a metric to gauge the reliability of each prediction, without resorting to validation using the mechanism-based model.

‘The wisdom of crowds’ refers to the phenomenon in which the collective knowledge of a community is greater than the knowledge of any individual^40^. To this end, we developed a ‘voting’ protocol, which relies on the training of several neural networks in parallel using the same set of training data. Even though these networks have the same architecture, the training process has an intrinsically stochastic component. Each network creates a map of virtual neurons and assigns random numerical values, or “weights”, to connections between them during the initialization process. If the network does not accurately predict a particular pattern, it will back-propagate the gradient of the error to each neuron and the weights would be updated to minimize the error in a new prediction. Even though the same rule is applied, each neuron is updated independently. With same training data, same architecture, the probability of getting exactly the same parameterized neural network is essentially zero. That is, each trained neural network is unique in terms of the parameterization of the network connections. Figure S6A illustrated the differences of trainable variables (weights, bias) between two trained neural networks. Despite the difference in parameterization, the different neural networks trained from the same data *overall* make similar predictions. We reasoned that this similarity can serve as the metric of the accuracy of the prediction. In particular, for a certain input parameter set, if all trained networks give very similar predictions, it is likely that these predictions are overall accurate. In contrast, for another input parameter set, if predictions from different networks diverge from each other, this divergence would suggest some or all of these predictions are not reliable. Given this reasoning, we could expect a positive correlation between the reliability of the prediction (in comparison to the correct prediction generated by the mechanism-based model) and the consistency between predictions generated by different neural networks.

To test this notion, we trained four neural networks. For each parameter combination in the testing set, we calculated the divergence between predictions by different neural networks, by using the MSE. The final prediction is the one with the least average MSE between all other predictions. We then calculated the divergence between the ensemble prediction and the correct profile. Indeed, the accuracy of the ensemble prediction is positively correlated with the consistency between different neural networks (Figure 4B, Figure S6B). That is, if predictions by neural networks exhibit high consensus, the errors in the prediction are also low. Also, a side benefit of using multiple neural networks is that the ensemble prediction is in general more reliable than one by a single neural network. The average MSE over the test dataset reduced from 0.0118 to 0.0066. This improved accuracy is expected and is the typical use of ensemble method^41^. The different predictors can be trained in parallel, via different GPU cores or even different servers on a computer cluster. Similarly, predictions can be made in parallel. This scalable ability makes it a powerful tool and may not affect the total time needed for training and for predicting.

**Figure 4.**
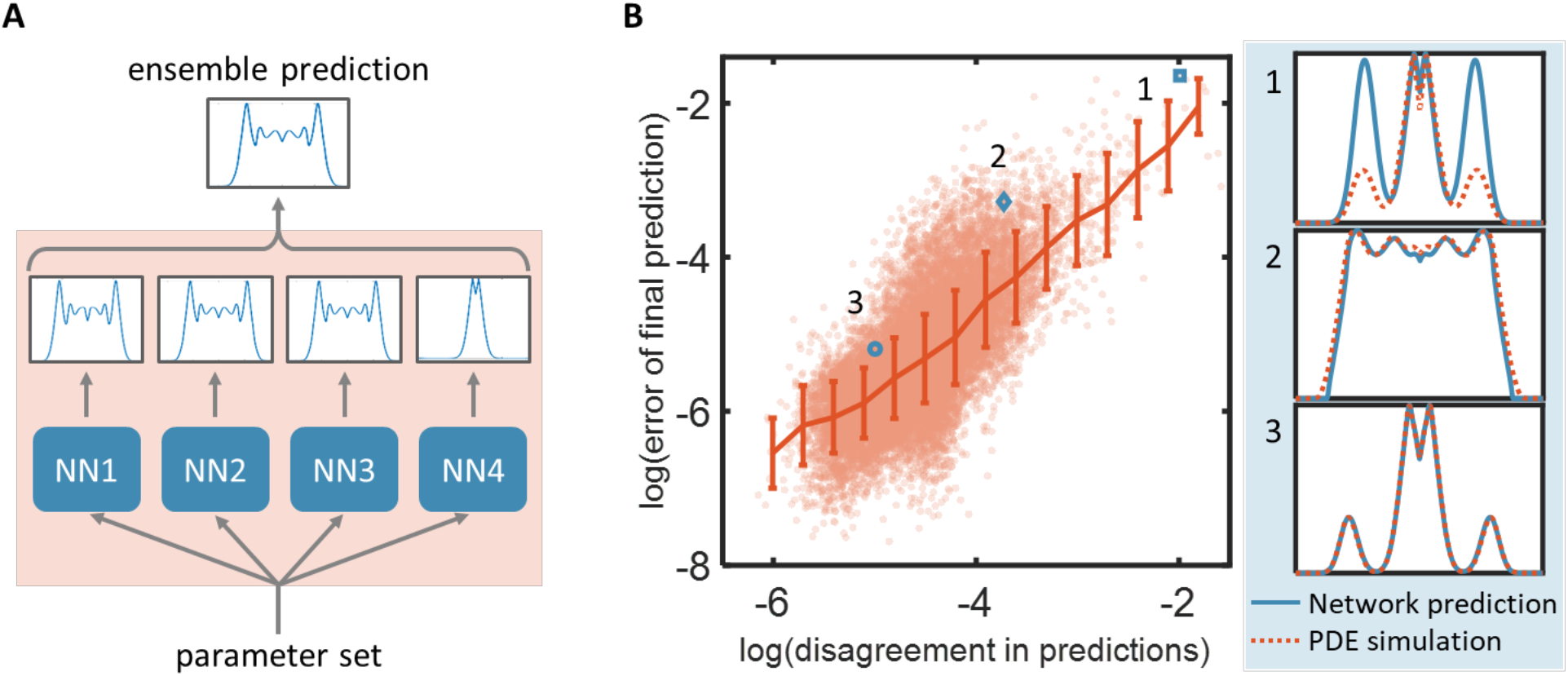
Ensemble predictions enable self-contained evaluation of the prediction accuracy. **A. Schematic plot of ensemble prediction.** With each new parameter set (different combinations of parameters), we have several neural networks to predict the distribution independently. Though these networks have the same architecture, the training process has an intrinsically stochastic component due to random initialization and backpropagation. There might be multiple solutions for a certain output to be reached due to non-linearity. Despite the difference in parameterization of trainable variables (Figure S6A), the different neural networks trained from the same data overall make similar predictions. Based on these independent predictions, we can get a finalized prediction using a ‘voting” algorithm (Details described in Methods). **B. The disagreement in predictions (DP) by neural networks trained in parallel is positively correlated with the error in prediction (EP).** We calculated the disagreement in predictions (averaged MSE between predictions from different neural networks) and error of final prediction (MSE between final ensemble prediction and PDE simulation) for all samples in test data set (red dots). We then divide all data into 15 equally-spaced bins and calculated the mean and variance for each bin (red line). The positive correlation suggests that the consensus between neural networks represents a self-contained metric for the reliability of each prediction. We showed three sample predictions with different degrees of accuracy: sample 1: DP=0.14, EP=0.19; sample 2: DP=0.024, EP=0.038; sample 3: DP=0.0068, EP=0.0056.

We screened through 10^8^ combination of parametric sets using the ensemble prediction method, where we discarded predictions with disagreement in predictions larger than 0.1. These neural network predictions reveal the general criterion for making complex patterns. For example, generation of 3-ring patterns requires a large domain radius (R), large synthesis rate of T7RNAP (*α_T_*), small synthesis rate of T7 lysozyme (*α_L_*), small half activation constant of T7RNAP (*K_T_*), small half activation distance for gene expression (*K_φ_*) (Figure 5A). Based on the analysis above, we want to further identify the correlation between *K_T_* and *α_C_, K_T_* and *α_T_, D* and *α_C_*. For each of the screening, we vary two parameters of interest and fixed the rest to identify the relationship between parameters required to generate 3-ring patterns. We collect 10^7^ instances and discard predictions with disagreement between ensemble predictions larger than 0.1(Figure 5B, Figure S7). We found that if the growth rate on agar (*α_C_*) is large, the domain radius (*D*) can be reduced. Additionally, there is a negative relationship between Cell growth rate on agar (*α_C_*) and half activation constant of T7RNAP (*K_T_*) (If approximating that they are inversely proportional, we can get the fitting with *R^2^* = 0.94. Figure S7A). We also found a linear correlation between half activation constant of T7RNAP (*K_T_*), and synthesis rate of T7RNAP (*α_T_*) in order to generate 3-ring patterns (*R^2^* = 0.996, Figure S7B). A key advantage is that machine-learning methods can sift through volumes of data to find patterns that would be missed otherwise. This provides significant insight in our experiments to find conditions that allow the formation of multiple rings, which could not be done using traditional simulation methods.

**Figure 5.**
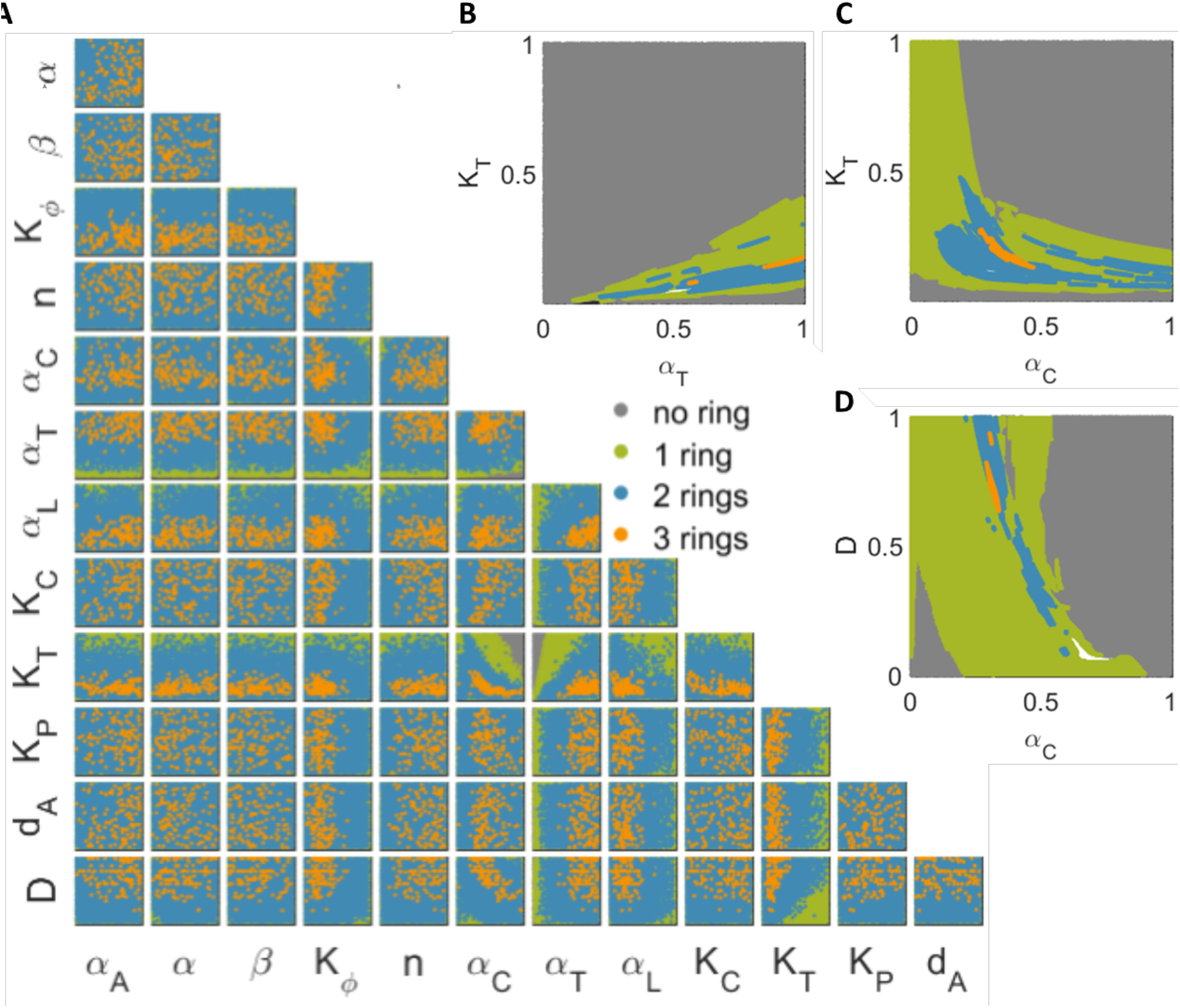
Neural network predictions enable comprehensive exploration of pattern formation dynamics in the model system. **A. Ensemble of deep neural networks enables screening through a vast parametric space.** The parametric space consists of 13 parameters that can vary uniformly in the provided ranges (Table S1). For each instance, we randomly generate all the varying parameters and use neural network to predict the peak and distributional values for that instance. We collect 10^8^ instances and discard predictions with disagreement between ensemble predictions larger than 0.1. We then project all the instances on all the possible 2D parameter plane. The majority of the instances generate patterns with no ring (grey), and they are distributed all over the projected parametric planes. Due to the huge number of instances, the parametric distribution of no ring (grey), 1-ring (green), 2-rings (blue) patterns on the projected 2D planes overlap on top of each other. From the distribution of neural network predicted 3-ring patterns (orange) over all the possible 2D parameter planes, The critical constraints to generate 3-ring patterns were revealed: large domain radius (*D*), large synthesis rate of T7RNAP (*α_T_*), small synthesis rate of T7 lysozyme (*α_L_*), small half activation constant of T7RNAP (*K_T_*), small half activation distance for gene expression (*K_φ_*). We also find possible correlations between *K_T_* and *α_C_* (cell growth rate on agar), *K_T_* and *α_T_*, *D* and *α_C_*. **B-D. The vast amount of neural network predictions greatly facilitates the evaluation the objective function of interest (generation of 3-ring patterns).** Based on the analysis above, we want to further identify the correlation between *K_T_* and *α_C_*, *K_T_* and *α_T_*, *D* and *α_C_*. For each of the screening, we vary two parameters of interest and fixed the rest. We collect 10^7^ instances and discard predictions with disagreement between ensemble predictions larger than 0.1. We found in order to generate 3-ring patterns, there is a negative correlation between *D* and *α_C_*, a negative correlation between *K_T_* and *α_C_*. We also found a positive linear correlation between *K_T_* and *α_T_*. (B) *α_A_* = 0.5, α = 0.5, β = 0.5, *K_ø_* = 0.3, *n* = 0.5, *α_T_* = 0.8, *α_L_* = 0.3, *K_C_* = 0.5, *K_P_* = 0.5, *d_A_* = 0.5, *K_T_* = 0.3. (C) *α_A_* = 0.5, α = 0.5, β = 0.5, *K_Ø_* = 0.3, *n* = 0.5, *α_T_* = 0.8, *α_L_* = 0.3, *K_C_* = 0.5, *K_P_* = 0.5, *d_A_* = 0.5, *D* = 1.0. (D) *α_A_* = 0.5, α = 0.5, β = 0.5, *K_ø_* = 0.3, *n* = 0.5, *α_C_* = 0.5, *α_L_* = 0.3, *K_C_* = 0.5, *K_P_* = 0.5, *d_A_* = 0.5, *D* = 1.0.

### The neural network predicts probability density functions from a stochastic model with high accuracy

Our framework is applicable to any dynamic model that generates a continuous series of each output. To illustrate this point, we apply the framework to the emulation of a stochastic model of the MYC/E2F pathway^42,43^ (Figure S7A). This model consists of 10 stochastic differential equations and 24 trainable parameters (see Methods and supplementary information). For each parameter set, repeated simulations lead to generation of the distribution of the levels of each molecule. With a sufficiently large number of simulations, this distribution converges to an approximately continuous curve (Figure S7B). As such, establishing the mapping between the parameter set and the corresponding output distribution is an identical problem as prediction of the spatial patterns. Again, the trained neural network exhibits high accuracy in predicting the distributions of different molecules in the model (Figure S8).

## Discussion

Our results demonstrate the tremendous potential of deep learning in overcoming the computational bottleneck faced by many mechanistic-based models. The key to the massive acceleration in predictions is to bypass the generation of fine details of system dynamics but instead focus on an empirical mapping between input parameters to system outputs of interest using NNs. The massive acceleration enables extensive exploration of the system dynamics that is impossible by solely dependent on the mechanistic model (Figure 5). Depending on the application context, this capability can facilitate the engineering of gene circuits or the optimization of experimental conditions to achieve specific target functions (e.g., generation of multiple rings from our circuit), or to elucidate how a biological system responds to environmental perturbations (e.g., drug treatments).

A major innovation of our approach is the combined use of the mechanistic model and the neural network (Figure 1). The mechanistic model is used as a stepping stone for the latter by providing a sufficient data set for training and testing. This training set is extremely small compared with the possible parameter space. Given the relatively small training set, the remarkable performance of the NN suggests that, for the models we tested, the landscape of the system outputs in the parametric space is sufficiently smooth. If so, a small training set is sufficient to reliably map the output landscape for the much broader parametric space. However, the NN does occasionally fail. We found that the NN tends to make more mistakes in clustered regions (e.g., the blank regions in Figure 5B, where data have been deleted due to large disagreement in predictions). As such, we suspect that the NN is more likely to fail when the system output is highly sensitive to local parameter variations, i.e., where the output landscape is rugged in the parametric space. Such a limitation could be alleviated by increasing the size of the training set. Alternatively, one could generate the training set according to local system sensitivity, such that training data are generated more densely in regions of high local sensitivity.

Our approach is generally applicable as long as each input parameter set generates a unique output (but the same output can correspond to different input parameter sets). This constraint is implied in both of our examples. In the pattern-formation circuit, each parameter combination can generate a unique final pattern. For the stochastic model, different runs of the model will generate different levels for each molecular species (for the same parameter set). However, the distribution of these levels for a sufficiently large number of simulations is approximately deterministic – it will be deterministic for an infinite number of simulations. Therefore, each parameter set in the stochastic model leads to a unique distribution for each molecule. This constraint is satisfied in vast majority of dynamical models of biological systems, where the output can be a time series, a spatial distribution, or distribution of molecules from ensemble simulations. As such, the general framework (Figure 1) is applicable to all these models. However, the benefit of the framework depends on the specific model of interest. In particular, our approach is most useful when the generation of the initial data set is nontrivial but manageable. While doable, our approach will not gain much by emulating simple ODE models, which can be solved quickly. Conversely, if a model is so complex, such that, even the generation of sufficient training data could be computationally prohibitive, an alternative integration of the mechanistic model and deep learning is necessary to speed up the training process

## Methods

### Modeling of pattern formation in Escherichia coli programed by a synthetic gene

The circuit consists of a mutant T7 RNA polymerase (T7RNAP) that activates its own gene expression and the expression of LuxR and LuxI. LuxI synthesizes an acyl-homoserine lactone (AHL) which can induce expression of T7 lysozyme upon binding and activating LuxR. Lysozyme inhibits T7RNAP and its transcription by forming a stable complex with it^44^. CFP and mCherry fluorescent proteins are used to report the circuit dynamics since they are co-expressed with T7RNAP and lysozyme respectively.

The PDE model used in the current study corresponds to the hydrodynamic limit of the stochastic agent-based model from Payne et al.^33^. Because the air pocket between glass plate and dense agar is only 20 μm high, the system was modeled in two spatial dimensions and neglect vertical variations in gene expression profiles^32^. Although the PDE formulation is computationally less expensive to solve numerically than the stochastic agent-based model and better facilitates development of mechanistic insights into the patterning dynamics, it still needs a lot of computational power when extensive parameter search is needed.

The circuit dynamics can be described by the following PDEs:

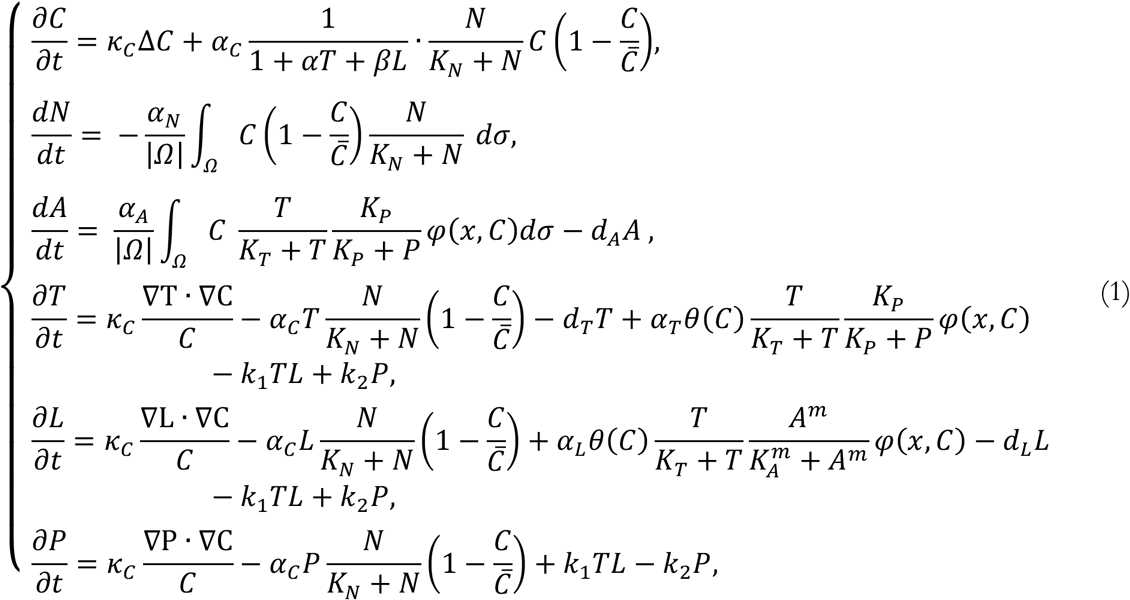

where *C*(*t, x*) is the cell density; *N*(*t*) is the nutrient concentration; *A*(*t*) is the AHL concentration; *T*(*t,x*), *L*(*t,x*), *P*(*t,x*) are cellular T7RNAP, lysozyme and the T7-lysozyme complex density respectively. See Table S1 for description of all model parameters.

### Modeling of MYC/E2F pathway in cell-cycle progression

The SDE model use in this study is the system describing the MYC/E2F pathway in cell-cycle progression^42,43,45^ (Figure S7A). Upon growth factor stimulation, increases in MYC lead to activation of E2F-regulated genes through two routes. First, MYC regulates expression of *Cyclin D*, which serves as the regulatory components of kinases that phosphorylate pocket proteins and disrupt their inhibitory activity. Second, MYC facilitates transcriptional induction of activator E2Fs, which activate the transcription of genes required for S phase. Expression of activator E2Fs is reinforced by two positive feedback loops. First, activator E2Fs can directly binds to their own regulatory sequences to help maintain an active transcription state, second, activator E2Fs transcriptionally upregulate CYCE, which stimulates additional phosphorylation of pocket proteins and prevents them from sequencing activator E2Fs.

To capture stochastic aspects of the Rb-E2F signaling pathway, we adopted the Chemical Langevin Formulation (CLF)^46^. We adjusted the units of the molecule concentrations and the parameters so that the molecules are expressed in molecular numbers.

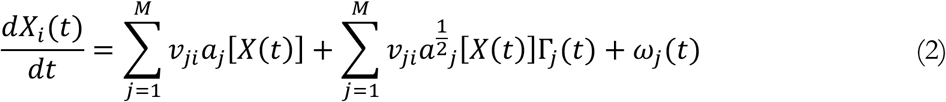

Where *X_i_*(*t*) represents the number of molecules of a molecular species I (i=1,…, N) at time t, and *X*(*t*) = (*X*_1_(*t*),…,*X_N_*(*t*)) is the state of the entire system at time t. The mean molecule number for E2F would be approximately 1,000. *X*(*t*) evolves over time at the rate of *a_j_*[*X*(*t*)] (j= 1,…, M), and the corresponding change in the number of individual molecules is described in *ν_ij_*. Γ_*j*_(*t*) and *ω_j_*(*t*) are temporally uncorrelated, statistically independent Gaussian noises. This formulation retains the deterministic framework (the first term), and intrinsic noise (reaction-dependent) and extrinsic noise (reaction-independent). The concentration units in the deterministic model were converted to molecule numbers, so that the mean molecule number for E2F would be approximately 1,000. We assumed a mean of 0 and variance of 5 for Γ_*j*_(*t*), and a mean of 0 and variance of 50 for *ω_j_*(*t*). The resulting stochastic differential equations (SDEs) were implemented and solved in Matlab. Serum concentration is fixed at [S] = 1%.

Twenty-four parameters of the SDE model are generated randomly at the range provided at Supplementary table S5. The range covers almost all the possible rates that can be found in vivo. For each of the generated combination of parameters, sample 10^4^ stochastic simulations and collect the final values of all 10 variables for each of the simulation. Each of the variables are discretized into 1,000 intervals. Create a kernel distribution object by fitting it to the data. Then by using Matlab function *pdf()* to get the probability density function of the distribution object, evaluated at the values in each of the discretized interval. (Matlab code can be found on GitHub, https://github.com/ShangyingWang/SDEmodel.git).

### Network performance

Mathematical equations for many biological applications can be described using diffusion-reaction equation: 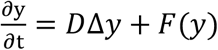, where y ∈ *R^m^* represents a group of physical or biological species, D ∈ *R^m×m^* is the diffusion constant matrix, Δy is the Laplacian associated with the diffusion of the species y, and F(y) describes the chemical or biological reactions. When the diffusion term is discretized using finite difference methods, the reaction diffusion system is first reduced to a series calculation of ODEs with the following form *y_t_* = *Ay* + *f*(*y*), where Ay is a finite difference approximation of *D*Δ*y*. Let *N* denote the number of spatial grid points for the approximation of Δ*y*, then y(t) ∈ *R^N×m^* and A is a (*N × m*) × (*N × m*) matrix and this need to be solved at every time step. The calculations on each grid and at each time step are often sequential processes and cannot be fastened using parallel computing. Because the size of the nonlinear system linearly depends on the spatial resolution N, using a standard nonlinear solver for such system is usually very expensive (computational complexity with an order of *N*^2^)^47^. The runtime of mechanism-based model varies, but usually needs hours or even days.

Most of the calculation of neural network are additions and multiplications and it also capable of running hundreds of thousands of matrix operations in a single clock cycle. Assuming each instance of numerical simulation needs 5 minutes and neural network prediction needs 0.01 seconds, our approach expedites the calculation 30,000 times.

### Data preparation, training and prediction

We used the Duke SLURM computing platform to simulate mechanism-based models and preparing data for training neural networks. We used Google cloud machine learning platform for hyper-parameter tuning. We use python 3.5 platform and implement TensorFlow 2.0 for neural network design and trainings/validations/tests. Source codes are available on github.com (https://github.com/ShangyingWang/pattern_prediction).

Machine learning algorithms do not perform well when the input numerical attributes have very different scales. During data preprocessing, we use min-max scaling to normalize all the input parameters to be within the region 0 to 1. We also extract the peak value from the distribution and log the peak value. We use LSTM network for prediction of the normalized distribution. Specifically, we divided the space along radius axis to 501 points. And each point is associated with an LSTM module (see below) for prediction.

### LSTM networks

Most of the neural networks are feedforward neural networks, where the information flows from the input layer to the output layer. A recurrent neural network (RNN) has connections pointing backwards. It will send the predicted output back to itself. An RNN when unrolled can be seen as a deep feed-forward neural network (Figure S2A). RNN is often used to predict time series data, such as stock prices. In autonomous driving systems, it can anticipate car trajectories and help avoid accidents.

However, the ordinary RNN cannot be used on long sequence data. The memory of the first inputs gradually fades away due to the transformations that the data goes through when traversing an RNN, some information is lost after each step. After a while, the RNN state contains virtually no trace of the first inputs^48^. To solve this problem, various types of cells with long-term memory have been introduced and the most successful/popular one is the LSTM network. Figure S2B showed the architecture of an LSTM cell. An internal recurrence (a self-loop) is added on top of the outer recurrence of the RNN. This self-loop is responsible for memorizing long-term dependencies (See supplementary information for more details).

### Network structure

Figure S2D demonstrates the structure of the employed Deep LSTM network, which consists of an input layer with inputs to be the parameters of mechanism-based model, a fully connected layer, LSTM arrays, and two output layers, one for predicting peak values of distributions, one for predicting the normalized distributions. First, the parameters of differential equations are connected to the neural network through a fully connected layer. Fully connected layer means all the inputs are connected to all the neurons in that layer. The activation function is **ELU** (**E**xponential **L**inear **U**nit) and the connection weight is initialized randomly using He initialization method^49^. It then connected to another fully connected layer with 1 neuron for peak value prediction. The output of the first fully connected layer is also connected to a sequence of LSTM modules for predicting distributions. We use Adam optimization algorithm to adaptive moment estimation and gradient clipping to prevent exploding gradients. (See supplementary information for more details).

### Network optimization

We used both cross entropy and MSE for calculating the cost function. Cross entropy originated from information theory. If the prediction is exactly the same as the pattern from simulation, cross entropy will just be equal to the entropy of the pattern from simulation. But if the prediction has some deviation, cross entropy will be greater by an amount called the Kullback-Leibler divergence (KLD).

The cross entropy between two distributions p and q is defined as H(p, q) = −Σ_*s*_*p*(*s*) log q(*s*). In our study, we found using either of the error function did not alter the accuracy of our network and the analysis.

### Ensemble prediction

To make a prediction for a new instance, we need to aggregate all the predictions from all predictors. The aggregation function is typically the statistical mode (i.e., the most frequent predictions) for classification problems and the average for regression problems. Previously, there is no study on what the aggregation function shall be for distributional predictions. Here, we proposed that for distributional predictions, similarity score between different predictions are calculated. The similarity score can be MSE, KL divergence, R^2^, or other similarity function between two distributions. In this paper, we choose MSE for calculating the similarity score. Each predictor is associated with a score based on the average of the MSE of its prediction in comparison with all other predictions. The final prediction will be the one with the minimal score.

In many cases, an aggregated answer from a group of people is often better than one person’s answer, which is called wisdom of the crowd^40^. Similarly, aggregating predictions of a group of predictors will often get better predictions than with only one predictor. A group of predictors is called an ensemble and this algorithm is called an ensemble method^41^.

Predictions from an ensemble of neural networks has lower error than predictions from a single neural network in predicting distributional data (Table S4). Also, the disagreement in prediction between predictors is positively associated with errors in predictions for test data set (Figure 4B).

Disagreement in prediction can be used as an estimate of the errors in predictions. In this way, we can rank our predictions with different confident levels.

## Acknowledgements

We thank Zhuojun Dai, Tatyana Sysoeva, Meidi Wang and Fred Huang for discussions and comments; Shiyuan Mei, Esther Brown and Yuanchi Ha for help editing the manuscript. Duke Compute Cluster (DCC) for assistance with high-performance computation solutions. This study was partially supported by the Office of Naval Research (N00014-12-1-0631), National Science Foundation (L.Y.) and National Institutes of Health (L.Y.: 1R01-GM098642; MDR: R01-GM096190).

