## supplemental file for "Massive computational acceleration by using neural networks to emulate mechanism-based biological models"

I. **Supplemental PDE model descriptions**
II. **Supplemental SDE model descriptions**
III. **Supplemental deep learning methods**
IV. **Supplemental tables**
V. **Supplemental figures and figure legends**
VI. **Supplemental references**

### I.Supplemental PDE model descriptions

#### Model development

The pattern formation circuit is from Payne, et al.[1]. The circuit consists of a mutant T7 RNA polymerase (T7RNAP) that activates its own expression as well as the expression of LuxR and LuxI. Once activated by T7RNAP, LuxI mediates synthesis of an acyl-homoserine lactone (AHL), which can diffuse across the cell membrane. When the global AHL concentration surpasses a threshold, intracellular AHL binds to LuxR to activate the synthesis of T7RNAP lysozyme. Lysozyme then binds to the T7RNAP and forms a complex, therefore inhibiting the T7RNAP binding to the T7RNAP promoter. This complex also inhibits T7RNAP transcription. CFP and mCherry fluorescent proteins are used to report the circuit dynamics since they are co-expressed with T7RNAP and lysozyme respectively (Figure S1).

The gene circuit dynamics can be described using the following partial differential equations (PDEs), which describe the cell growth, colony expansion, nutrient and AHL diffusion, intracellular circuit dynamics, as well as signaling and transport. (Parameters are described in Table S1). This PDE model corresponds to the hydrodynamic limit of the stochastic agent-based model from Payne et al.[1]. Because the air pocket between the glass plate and dense agar is only 20 μm high, the system was modeled in two spatial dimensions and vertical variations in gene expression profiles were neglected. Although the PDE formulation is computationally less expensive to solve numerically than the stochastic agent-based model and better facilitates the development of mechanistic insights into the patterning dynamics, it still needs a lot of computational power when extensive parameter search is needed.

$$
\begin{cases}
\dfrac{\partial C}{\partial t} = \kappa_C \Delta C + \alpha_C \dfrac{1}{1+\alpha T+\beta L} \cdot \dfrac{N}{K_N+N} C\left(1-\dfrac{C}{\bar{C}}\right), \\
\dfrac{dN}{dt} = -\dfrac{\alpha_N}{|\Omega|}\displaystyle\int_\Omega C\left(1-\dfrac{C}{\bar{C}}\right)\dfrac{N}{K_N+N}\, d\sigma, \\
\dfrac{dA}{dt} = \dfrac{\alpha_A}{|\Omega|}\displaystyle\int_\Omega C \dfrac{T}{K_T+T}\dfrac{K_P}{K_P+P}\varphi(x,C) d\sigma - d_A A, \\
\dfrac{\partial L}{\partial t} = \kappa_C \dfrac{\nabla \mathrm{L}\cdot\nabla \mathrm{C}}{C} - \alpha_C L \dfrac{1}{1+\alpha T+\beta L}\dfrac{N}{K_N+N}\left(1-\dfrac{C}{\bar{C}}\right) - d_L L + \alpha_L \dfrac{T}{K_T+T}\dfrac{A^m}{K_A^m+A^m}\varphi(x,C) - k_1 TL + k_2 P, \\
\dfrac{\partial T}{\partial t} = \kappa_C \dfrac{\nabla \mathrm{T}\cdot\nabla \mathrm{C}}{C} - \alpha_C T \dfrac{1}{1+\alpha T+\beta L}\dfrac{N}{K_N+N}\left(1-\dfrac{C}{\bar{C}}\right) - d_T T + \alpha_T \dfrac{T}{K_T+T}\dfrac{K_P}{K_P+P}\varphi(x,C) - k_1 TL + k_2 P, \\
\dfrac{\partial P}{\partial t} = \kappa_C \dfrac{\nabla \mathrm{P}\cdot\nabla \mathrm{C}}{C} - \alpha_C P \dfrac{1}{1+\alpha T+\beta L}\dfrac{N}{K_N+N}\left(1-\dfrac{C}{\bar{C}}\right) + k_1 TL - k_2 P, \\
\dfrac{\partial \psi_R}{\partial t} = \kappa_C \dfrac{\nabla\psi_R\cdot\nabla C}{C} - \alpha_C \psi_R \dfrac{1}{1+\alpha T+\beta L}\dfrac{N}{K_N+N}\left(1-\dfrac{C}{\bar{C}}\right) + \alpha_L \dfrac{T}{K_T+T}\dfrac{A^m}{K_A^m+A^m}\varphi(x,C), \\
\dfrac{\partial \psi_C}{\partial t} = \kappa_C \dfrac{\nabla\psi_C\cdot\nabla C}{C} - \alpha_C \psi_C \dfrac{1}{1+\alpha T+\beta L}\dfrac{N}{K_N+N}\left(1-\dfrac{C}{\bar{C}}\right) + \alpha_T \dfrac{T}{K_T+T}\dfrac{K_P}{K_P+P}\varphi(x,C).
\end{cases} \quad (1)
$$

$C(t,x)$ is the cell density; $N(t)$ is the nutrient concentration; $A(t)$ is the AHL concentration; $L(t,x), T(t,x), P(t,x)$ are cellular lysozyme, T7RNAP and the T7-lysozyme complex density respectively; $\psi_R(t,x)$ and $\psi_C(t,x)$ are mCherry and CFP, which are co-expressed with lysozyme and T7RNAP, respectively, and act as reporters in experiments. These are added in order to allow for a direct comparison between model and experiment;

The following assumptions were made in deriving these equations:

1. Cells are assumed to perform an unbiased random walk; their movement is modeled as diffusion[2-4]. We considered "diffusion" as an approximation of the observed colony expansion, so that cell movement can be described by a single lumped parameter. Intracellular components are modeled with passive-tracer equations (see derivation below).

2. Cell growth is modeled by a logistic term, along with a Monod function. The Monod function is to account for the contribution of nutrient to overall colony growth. The nutrient here refers to one or more limiting factors that constrain growth. The logistic term accounts for the limit of cell growth in a particular location. This carrying capacity is unlikely limited by nutrient availability. Instead, it is limited by the spatial confinement imposed by our device, which is the colony height confined to be ~20 $\mu m$ between the coverslip and the agar surface.

3. Fast diffusion of AHL and nutrient.

4. Gene expression capacity:

$$\varphi(x, C) = \begin{cases} \dfrac{K_\varphi^n}{K_\varphi^n + \left(R_\varphi - x\right)^n}, & x \le R_\varphi \\ 1, & x > R_\varphi \end{cases} \tag{2}$$

where $R_\varphi$ is defined as the distance between the colony center and the location where cell density is 95% of the carrying capacity.

5. Assume that $L, T$ and $P$ are at equilibrium due to the reversible first-order kinetics of T7RNAP bind with T7 lysozyme to form T7-lysozyme complex is fast[5].

$$P = \frac{k_1}{k_2} TL \tag{3}$$

**Non-dimensionalization of the model**

First, we rescaled the time and space variables as

$$\hat{t} = \alpha_C t, \quad \hat{x} = \frac{x}{\mathcal{L}}, \tag{4}$$

where $\mathcal{L}$ is a length scale to be chosen later.

We next rescaled the state variables,

$$\hat{C} = \frac{C}{\bar{C}}, \ \hat{N} = \frac{N}{N_0}, \ \hat{A} = \frac{A}{K_A}, \ \hat{L} = \frac{d_L}{\alpha_L} L, \ \hat{T} = \frac{T}{K_T}, \ \hat{P} = \frac{P}{K_P}, \ \widehat{\psi_R} = \frac{\psi_R}{\alpha_L}, \ \widehat{\psi_C} = \frac{\psi_C}{\alpha_T}. \tag{5}$$

Then we defined some new parameters for simplicity,

$$\hat{\alpha} = \alpha K_T, \ \hat{\beta} = \frac{\alpha_L}{d_L} \beta. \tag{6}$$

With these dimensionless variables, and by defining $\hat{\varphi}\left(\hat{x}, \hat{C}\right) = \varphi(x, C)$, we can rewrite the model equations in a dimensionless form. Introducing the parameter groups $G_i, (i = 1, \ldots, 12)$ (see Table S2), the non-dimensioned equations become:

$$
\begin{cases}
\frac{\partial \hat{C}}{\partial t} = G_1 \Delta \hat{C} + \frac{1}{1+\hat{\alpha}\hat{T}+\hat{\beta}\hat{L}} \hat{C}(1-\hat{C}) \frac{\hat{N}}{G_2+\hat{N}} , \\
\frac{d\hat{N}}{dt} = -G_3 \int_{\Omega} \hat{C}(1-\hat{C}) \frac{\hat{N}}{G_2+\hat{N}} d\sigma , \\
\frac{d\hat{A}}{dt} = G_4 \int_{\Omega} \hat{C} \frac{\hat{T}}{1+\hat{T}} \frac{1}{1+\hat{P}} \varphi(\hat{x},\hat{C}) d\sigma - G_5 \hat{A} , \\
\frac{\partial \hat{L}}{\partial t} = G_1 \frac{\nabla \hat{L} \cdot \nabla \hat{C}}{\hat{C}} - \hat{L} \frac{1}{1+\hat{\alpha}\hat{T}+\hat{\beta}\hat{L}} \frac{\hat{N}}{G_2+\hat{N}} (1-\hat{C}) - G_6 \hat{L} + G_7 \frac{\hat{T}}{1+\hat{T}} \frac{\hat{A}^m}{1+\hat{A}^m} \varphi(\hat{x},\hat{C}) , \\
\frac{\partial \hat{T}}{\partial t} = G_1 \frac{\nabla \hat{T} \cdot \nabla \hat{C}}{\hat{C}} - \hat{T} \frac{1}{1+\hat{\alpha}\hat{T}+\hat{\beta}\hat{L}} \frac{\hat{N}}{G_2+\hat{N}} (1-\hat{C}) - G_8 \hat{T} + G_9 \frac{\hat{T}}{1+\hat{T}} \frac{1}{1+\hat{P}} \varphi(\hat{x},\hat{C}) , \\
\frac{\partial \hat{P}}{\partial t} = G_1 \frac{\nabla \hat{P} \cdot \nabla \hat{C}}{\hat{C}} - \hat{P} \frac{1}{1+\hat{\alpha}\hat{T}+\hat{\beta}\hat{L}} \frac{\hat{N}}{G_2+\hat{N}} (1-\hat{C}) , \\
\frac{\partial \widehat{\psi_R}}{\partial t} = G_1 \frac{\nabla \widehat{\psi_R} \cdot \nabla \hat{C}}{\hat{C}} - \widehat{\psi_R} \frac{1}{1+\hat{\alpha}\hat{T}+\hat{\beta}\hat{L}} \frac{\hat{N}}{G_2+\hat{N}} (1-\hat{C}) + G_{mcherry} \frac{\hat{T}}{1+\hat{T}} \frac{\hat{A}^m}{1+\hat{A}^m} \varphi(\hat{x},\hat{C}) , \\
\frac{\partial \widehat{\psi_C}}{\partial t} = G_1 \frac{\nabla \widehat{\psi_C} \cdot \nabla \hat{C}}{\hat{C}} - \widehat{\psi_C} \frac{1}{1+\hat{\alpha}\hat{T}+\hat{\beta}\hat{L}} \frac{\hat{N}}{G_2+\hat{N}} (1-\hat{C}) + G_{CFP} \frac{\hat{T}}{1+\hat{T}} \frac{1}{1+\hat{P}} \varphi(\hat{x},\hat{C}) .
\end{cases}
\tag{7}
$$

The definition of parameter groups $G_i, (i = 1, \ldots, 12)$ can be found in Table S2. Equation 3 becomes

$$
\hat{P} = \frac{G_{10}}{G_{11}G_{12}} \hat{T}\hat{L} \tag{8}
$$

##### Numerical solver for the PDE model

To solve the model numerically in Matlab, we exploit the radial symmetry of the system and reduce it to a PDE in polar coordinates, only depending on one spatial variable, namely the radius $r \in [0, R]$. We combine the Matlab built-in Runge-Kutta solver ode45 with a second order centered finite difference scheme for discretization of the gradients. Due to the radial symmetry, we use the 1D distribution along the radius as the ground truth for training/testing the neural network (Figure S1B) without losing any information.

In addition, due to the assumption that $L, T$ and $P$ are at equilibrium, the *L-T-P* system is updated in each step by projecting it onto the manifold defined by $P = \frac{G_{10}}{G_{11}G_{12}} TL$. With this constraint, the concentrations are updated to $(L_1, T_1, P_1)$

$$
L_1 = \frac{1}{2}\Big(L_0 - G_{10}T_0 - G_{11} + \sqrt{(L_0 - G_{10}T_0 - G_{11})^2 + 4G_{11}(L_0 + G_{12}P_0)}\Big),
$$

$$P_1 = P_0 + \frac{1}{G_{12}}(L_0 - L_1),$$

$$T_1 = T_0 - \frac{1}{G_{10}}(L_0 - L_1).$$

Although the PDE model is computationally less expensive than the stochastic agent-based model[1], it still imposes a prohibitive barrier for practical applications while intensive parameter searching or estimation are needed, even when computer clusters are used.

##### Parameters screening and the execution of PDE model

Each dimensionless parameter (Table S1) is a combination of several parameters with units (Table S2). Rather than estimating dimensionless parameters directly, we search values of dimensional parameters in a realistic range, and then determine the corresponding dimensionless parameters. We have 13 dimensional parameters randomly picked from a predefined range, some due to lack of literature estimations/measurements; some can be tuned with varying pH, temperature, nutrient, agar density and other factors (marked bold in Table S1). Other parameters are fixed with specific values either from literature or from experiments.

#### II. Supplemental SDE model descriptions

##### Model description

The deterministic Ordinary differential equations (ODE) for the Myc-E2F system, developed in the previous work, served as the basis for the stochastic Rb-E2F model[6,7].

$$
\begin{cases}
\frac{d[MC]}{dt} = \frac{k_{MC}[S]}{K_S + [S]} - d_{MC}[\mathrm{MC}]\,, \\
\frac{d[EFm]}{dt} = \left(k_S \frac{[S]}{K_S + [S]} + k_{EFm}\frac{[MC]}{K_{MC} + [MC]}\frac{[EFp]}{K_{EF} + [EFp]} + \frac{k_b[MC]}{K_{MC} + [MC]}\right)\frac{K_R}{K_R + [MC]} - d_{EFm}[\mathrm{EFm}], \\
\frac{d[EFp]}{dt} = k_{EFp}[EFm]\frac{K_{MR}}{K_{MR} + [MR]} + \frac{k_{p1}[CD][RE]}{K_{CD} + [RE]} + \frac{k_{p2}[CE][RE]}{K_{CE} + [RE]} - k_{RE}[\mathrm{RB}][\mathrm{EFp}] \\
\qquad - (1 + K_{AFR}[\mathrm{AF}])d_{EFp}[\mathrm{EFp}] \\
\frac{d[CD]}{dt} = \frac{k_{CD}[MC]}{K_{MCCD} + [MC]} + \frac{k_{CDS}[S]}{K_S + [S]} - d_{CD}[\mathrm{CD}] \\
\frac{d[CE]}{dt} = \frac{k_{CE}[EFp]}{K_{EF} + [EFp]} - d_{CE}[\mathrm{CE}], \\
\frac{d[RB]}{dt} = k_{RB} + \frac{k_{DP}[RP]}{K_{RP} + [RP]} - k_{RE}[\mathrm{RB}][\mathrm{EFp}] - \frac{k_{p1}[CD][RB]}{K_{CD} + [RB]} - \frac{k_{p2}[CE][RB]}{K_{CE} + [RB]} - d_{RB}[\mathrm{RB}], \\
\frac{d[RP]}{dt} = \frac{k_{p1}[CD][RB]}{K_{CD} + [RB]} + \frac{k_{p2}[CE][RB]}{K_{CE} + [RB]} + \frac{k_{p1}[CD][RE]}{K_{CD} + [RE]} + \frac{k_{p2}[CE][RE]}{K_{CE} + [RE]} - \frac{k_{DP}[RP]}{K_{RP} + [RP]} - d_{RP}[\mathrm{RP}]\,, \\
\frac{d[RE]}{dt} = k_{RE}[\mathrm{RB}][\mathrm{EFp}] - \frac{k_{p1}[CD][RE]}{K_{CD} + [RE]} - \frac{k_{p2}[CE][RE]}{K_{CE} + [RE]} - d_{RE}[\mathrm{RE}]\,, \\
\frac{d[AF]}{dt} = k_{AFb} + k_{AFMC}\frac{[MC]}{K_{AFMC} + [MC]} + \frac{k_{AFEF}[EFp]}{K_{AFEF} + [EFp]} - d_{AF}[\mathrm{AF}]\,, \\
\frac{d[MR]}{dt} = k_{MRMC}\frac{[MC]}{K_{MRMC} + [MC]} + \frac{k_{MREF}[EFp]}{K_{MREF} + [EFp]} - d_{MR}[\mathrm{MR}]\,,
\end{cases}
\tag{9}
$$

where $[S]$ is the growth signals (e.g. serum); $[MC]$, $[EFm]$, $[EFp]$, $[CD]$, $[CE]$, $[RB]$, $[RP]$, $[AF]$, $[MR]$ are the concentrations of Myc, E2F mRNA, E2F protein, CycD, CycE, Rb and Phosphorylated Rb, ARF and miRNA. $[RE]$ is the concentration of RB-E2F complex.

Initial conditions:

[RB]=[RE]=[M]=[E]=[CD]=[CE]=[RP]=0μM.

Parameters are defined in Table S5.

The above is the deterministic ODE model of the system. To capture stochastic aspect of the Rb-E2F signaling pathway, we adopt the Chemical Langevin Formulation (CLF)[8]. We adjust the units of the molecule concentrations and the parameters so that the molecules are expressed in molecular numbers.

$$\frac{dX_i(t)}{dt} = \sum_{j=1}^{M} v_{ji} a_j[X(t)] + \sum_{j=1}^{M} v_{ji} a^{\frac{1}{2}}{}_j[X(t)]\Gamma_j(t) + \omega_j(t) \tag{10}$$

$X_i(t)$ represents the number of molecules of a molecular species I (i=1,…, N) at time t, and $X(t) = (X_1(t), \dots, X_N(t))$ is the state of the entire system at time t. The mean molecule number for E2F would be approximately 1,000. X(t) evolves over time at the rate of $a_j[X(t)]$ (j= 1, …, M), and the corresponding change in the number of individual molecules is described in $v_{ji}$. $\Gamma_j(t)$ and $\omega_j(t)$ are temporally uncorrelated, statistically independent Gaussian noises. This formulation retains the deterministic framework (the first term), and intrinsic noise (reaction-dependent) and extrinsic noise (reaction-independent). The concentration units in the deterministic model were converted to molecule numbers, so that the mean molecule number for E2F would be approximately 1,000. We assumed a mean of 0 and variance of 5 for $\Gamma_j(t)$, and a mean of 0 and variance of 50 for $\omega_j(t)$. The resulting stochastic differential equations (SDEs) were implemented and solved in Matlab. Serum concentration is fixed at $[\mathrm{S}] = 1\%$.

Twenty-four parameters of the SDE model are generated randomly (table S5). The range covers almost all the possible rates that can be found in vivo. For each of the generated combination of parameters, we sample $10^4$ stochastic simulations and collect the final values of all 10 variables. We split the values into 100 bins to construct a histogram for each variable. Since the large number of simulations, the histograms are almost continuous. We create a kernel distribution object by using MATLAB function fitdist(). Then we use Matlab function *pdf()* to get the probability density function of the distribution object, evaluated at the values in each of the discretized intervals (Each of the variables are discretized into 1,000 intervals for this model).

### III.Supplemental deep learning methods

Deep learning through the training of artificial neural networks has made immense contributions in various fields, such as computer vision[9-11], speech recognition[12-15], and beating the world champion at the game of Go[16-18]. This is a result of fast GPUs, high availability of data, and also the advancements of the algorithms for training deep neural networks. Over the last decade, deep learning is also becoming increasingly important for diverse biological researches[19-26]. Among all the applications, a predictive model was developed based on statistical associations among features of a given dataset. The learned model can then be used to predict desired outputs, such as binary responses (e.g., pathogenic or non-pathogenic, toxic or non-toxic), categorical labels (e.g., bacteria strains, stages of diseases), values (e.g., growth rate, drug doses) or sequences (e.g., time/spatial series, probability density functions).

#### Recurrent neural networks

Recurrent Neural networks (RNNs) are a family of deep neural networks for processing sequential data[27-29]. Different from a feedforward neural network, a recurrent neural network has connections pointing backward. It will send the predicted output back to itself. Figure S2A is a demonstration of a recurrent neuron (the simplest RNN, composed of only one neuron receiving inputs, producing outputs, and sending the outputs back to itself). At each sequential step (also called a frame), this recurrent neuron receives input $x_s$ as well as its own output from previous sequential step $y_{s-1}$). By unrolling the network against the sequential inputs, we can see that each member of the output is a function of the previous output, and is produced using the same update rule applied to the previous outputs, which results in the sharing of parameters through a very deep computational graph.

$$y_s = h(y_{s-1}; x_s; \theta) = h(h(y_{s-2}; x_{s-1}; \theta); x_s; \theta) = h(h(h(y_{s-3}; x_{s-2}; \theta); x_{s-1}; \theta); x_s; \theta)$$

Since the output of a recurrent neuron at step $s$ is a function of all the inputs from previous steps, it seems to have a form of memory. However, the ordinary RNN cannot be used on long-sequence data. The memory of the first inputs gradually fades away due to the transformations that the data goes through when traversing an RNN, some information is lost after each step. After a while, the RNN state contains virtually no trace of the first inputs[30]. To solve this problem, various types of cells with long-term memory have been introduced and the most successful/popular one is the LSTM network.

##### LSTM network

The LSTM network was proposed in 1997 by Sepp Hochreiter and Jurgen Schmidhuber[31], and it was gradually improved over the years by Alex Graves[32], Wojciech Zaremba[33], and many more. Figure S2B showed the architecture of an LSTM cell. An internal recurrence (a self-loop, shown in red) is added on top of the outer recurrence of the RNN (shown in orange). This self-loop is responsible for memorizing long-term dependencies[29]. LSTM also has more parameters and a system of gating units to control the flow of information. The state unit, which has the linear self-loop, is the most important component and its weight is controlled by a forget gate unit (The weight can be a value between 0 and 1 via a sigmoid unit).

In order to use LSTM network to predict the distribution, we need to discretize the distribution into a sequence of $n$ consecutive values ($n$=501 for the first example). Each value is associated with an LSTM module. So there are 501 LSTM modules in our deep LSTM network for predicting the synthetic patterns. For each of the LSTM module, the inputs consist both the outputs from fully connected layer and outputs of the previous $m$ neighboring LSTM modules ($m$=16 in Figure S2C demonstration). The output of each LSTM module ($LSTM_i$) is a single value corresponding to the $i$th value among the $n$ consecutive values.

Figure S2D demonstrates the structure of the employed deep LSTM network, which consists of an input layer with inputs to be the parameters of mechanism-based model, a fully connected layer (with $l$ nodes), LSTM arrays (consist of $n$ LSTM modules), and two output layers, one for predicting peak values of distributions, one for predicting the normalized distributions. First, the parameters of differential equations are connected to the neural network through a fully connected layer. Fully connected layer means all the inputs are connected to all the neurons in that layer. The activation function is **ELU** (**E**xponential **L**inear **U**nit) and the connection weight is initialized randomly using *He* initialization method[34]. It is then connected to another fully connected layer with 1 neuron for peak value prediction. The output of the first fully connected layer is also connected to a sequence of LSTM modules for predicting distributions. We use Adam optimization algorithm (the momentum decay hyperparameter $\beta_1 = 0.5$, the scaling decay hyperparameter $\beta_2 = 0.999$) to adaptive moment estimation and gradient clipping to prevent exploding gradients.

To predict the patterns from PDE model demonstrated in this paper, we use $l$=64, $n$=501, $m$=16. To predict the probability distribution from SDE model demonstrated in this paper, we use $l$=256, $n$=1000, $m$=64. Number of units is chosen to be equal to 256 for each LSTM module. The initial learning rate is $10^{-4}$.

The learning process itself refers to finding the optimal set of model parameters that translate the features in the input data into accurate predictions of the labels. The parameters are found through a series of back and forth steps (a.k.a. backpropagation), where parameters are estimated, the model performance is evaluated, errors are identified and corrected, and then the process repeats, until the model performance cannot be improved upon, which is assessed by the minimization of the model error. Once the optimal parameters are identified, the model can be used to make predictions using new data.

#### IV. Supplemental tables

**Table S1. Definitions and the values of parameters used in the PDE model**

- To generate the training/test datasets, the values of 13 parameters were randomly picked from prespecified ranges (bold) and other parameters were fixed.
- The values of parameters mentioned in the main text are normalized to be between 0 and 1 with the following formulation: normalized parameter value = (parameter value-min)/(max-min).

| Parameter | Description | Defined value or search range | Base Unit |
|---|---|---|---|
| $k_1$ | Combination rate of T-Lys complex[5] | 400 | $molecule^{-1}h^{-1}\cdot cell$ |
| $k_2$ | Dissociation rate of T-Lys complex[5] | 10800 | $h^{-1}$ |
| $k_D\ (= k_1/k_2)$ | Equilibrium association constant of T7-lysozyme complex[5] | 0.037 | $molecule^{-1}\cdot cell$ |
| $\boldsymbol{\kappa_C}$ | **Cellular diffusion coefficient (depend on agar density)[35]** | **0.001-0.005** | **$cm^2\cdot h^{-1}$** |
| $\boldsymbol{\alpha_C}$ | **Cell growth rate on agar** | **0.2-2** | **$h^{-1}$** |
| $\alpha_N$ | Nutrient depletion rate (Fit with experiments) | 155 | $molecule\cdot h^{-1}\cdot cell^{-1}$ |
| $\boldsymbol{\alpha_A}$ | **AHL synthesis rate[36]** | **20- $2.0\times10^5$** | **$molecule\cdot h^{-1}\cdot cell^{-1}$** |
| $\boldsymbol{\alpha_L}$ | **Synthesis rate of T7 lysozyme** | **90 - $9\times10^3$** | **$molecule\cdot h^{-1}\cdot cell^{-1}$** |
| $\boldsymbol{\alpha_T}$ | **Synthesis rate of T7RNAP** | **80 - $8\times10^3$** | **$molecule\cdot h^{-1}\cdot cell^{-1}$** |
| $\boldsymbol{d_A}$ | **AHL degradation rate[36]** | **0.05-2** | **$h^{-1}$** |
| $d_L$ | Degradation rate of T7 lysozyme[1] | 0.0144 | $h^{-1}$ |
| $d_T$ | Degradation rate of T7RNAP[1] | 0.3 | $h^{-1}$ |
| $K_A$ | Concentration threshold of AHL to half-maximum of the pLuxI promoter[37] | 20 | nM |
| $K_N$ | Half-saturation for nutrient uptake (Fit with experiments) | 20 | nM |
| $\boldsymbol{K_T}$ | **Half activation constant of T7RNAP** | **50 - $5\times10^3$** | **$molecule\cdot cell^{-1}$** |
| $\boldsymbol{K_P}$ | **Half inhibition of T-Lys complex** | **50 - $5\times10^3$** | **$molecule\cdot cell^{-1}$** |
| $\boldsymbol{K_\varphi}$ | **Half activation distance for gene expression** | **0 - 10** | **cm** |
| $\boldsymbol{\hat{\alpha}}$ | **Inhibition factor of T7RNAP on Growth** | **0 - 5** | |
| $\boldsymbol{\hat{\beta}}$ | **Inhibition factor of T7 lysozyme on Growth** | **0 - $2\times10^3$** | |
| $m$ | Hill coefficient of AHL mediated gene expression[1] | 2 | |

| | | | |
|---|---|---|---|
| $\boldsymbol{n}$ | **Hill coefficient for distance-dependent gene expression capacity** | **0 - 5** | |
| $\bar{C}$ | Cell carrying capacity (Fit with experiments) | $3\times10^5$ | cells·ml$^{-1}$ |
| $\mathcal{L}$ | Non-dimensionalized factor for space (Fit with experiments) | 0.18898 | cm |
| $N_0$ | Initial nutrient concentration (Fit with experiments) | 66.67 | nM |
| $\lvert\Omega\rvert_0$ | Normalization factor for domains (Fit with experiments) | $1.69\times 10^{-8}$ | cm$^2$ |
| **D**$(=\frac{\lvert\Omega\rvert}{\lvert\Omega\rvert_0})$ | **Non-dimensionalized domain radius** | **1.0-3.0** | |

**Table S2. Expressions and values of parameters in non-dimensional model**

| Non-dimensionalized parameter | Expression | Value |
|---|---|---|
| $G_1$ | $\frac{\kappa_C}{\alpha_C \mathcal{L}^2}$ | $28 \times \frac{\kappa_C}{\alpha_C}$ * |
| $G_2$ | $\frac{K_N}{N_0}$ | $0.3$ |
| $G_3$ | $\frac{\alpha_N \bar{C}}{\alpha_C N_0} \frac{\mathcal{L}^3}{\lvert\Omega\rvert} \frac{1}{10^{-4} cm}$ | $0.0046 \times \frac{1}{\alpha_C D^2}$ * |
| $G_4$ | $\frac{\alpha_A \bar{C}}{\alpha_C K_A} \frac{\mathcal{L}^3}{\lvert\Omega\rvert} \frac{1}{10^{-4} cm}$ | $9.95 \times 10^{-5} \times \frac{\alpha_A}{\alpha_C D^2}$ * |
| $G_5$ | $\frac{d_A}{\alpha_C}$ | $\frac{d_A}{\alpha_C}$ |
| $G_6$ | $\frac{d_L}{\alpha_C}$ | $\frac{0.0144}{\alpha_C}$ |
| $G_7$ | $\frac{d_L}{\alpha_C}$ | $\frac{0.0144}{\alpha_C}$ |
| $G_8$ | $\frac{d_T}{\alpha_C}$ | $\frac{0.3}{\alpha_C}$ |
| $G_9$ | $\frac{\alpha_T}{\alpha_C K_T}$ | $\frac{\alpha_T}{\alpha_C K_T}$ |
| $G_{10}$ | $\frac{K_T d_L}{\alpha_L}$ | $\frac{0.0144 \times K_T}{\alpha_L}$ |
| $G_{11}$ | $\frac{d_L}{\alpha_L k_D}$ | $\frac{0.3892}{\alpha_L}$ |
| $G_{12}$ | $\frac{K_P d_L}{\alpha_L}$ | $\frac{0.0144 \times K_P}{\alpha_L}$ |
| $G_{mCherry}$ | $\frac{1}{\alpha_C}$ | $\frac{1}{\alpha_C}$ |
| $G_{CFP}$ | $\frac{1}{\alpha_C}$ | $\frac{1}{\alpha_C}$ |

* * By comparing the experiment colony expansion with Fisher-KPP's traveling wave solution with wave speed [38], we can estimate that $\mathcal{L}$ =0.18898 cm=1889.8 μm.

**Table S3. Using the mechanism-based model to validate 3-ring patterns generated from LSTM networks.** We use an ensemble of trained deep LSTM networks to screen through the parameter space. It takes around 12 days to screen through $10^8$ combinations of parameter sets, which would need thousands of years if we could generate these by using PDE simulations. We find 1284 three-ring pattern distributions, including novel patterns not present in the training sets. We then use their parameter combinations as inputs to generate numerical simulations from the PDE model and compare the distributions generated from LSTM network and from numerical simulations, we find most of the distributions from numerical simulations are consistent with that from network predictions. The mean value of the MSEs between NN predicted distributions and PDE simulations is 0.079 and the standard deviation is 0.008. If setting the threshold of mean squared value (MSE) between distributions generated by the neural network and the distributions generated by numerical simulations to be 0.1, there are 1203 found distributions with MSE<0.1 and only 81 with MSE>0.1.

| Total | MSE<0.1 | MSE>0.1 |
|---|---|---|
| 1284 | 1203 | 81 |

**Table S4. MSE of the test dataset for the PDE model.** We calculate the MSEs between predicted distributions from network predictions and that from numerical simulations for test dataset. The network prediction can be from one neural network, or an ensemble of neural networks (4 ensembles are chosen for comparison in this table). Since the final prediction of an ensemble of neural networks is based on the disagreement of distributional values, we can see ensemble neural networks have better accuracy on predicting distributional value than, and almost the same accuracy on predicting peak value as, single neural network.

| | | MSE | | |
|---|---|---|---|---|
| | | no ring | 1 ring | 2 rings and more |
| single NN | peak value | 0.35 | 0.19 | 0.14 |
| | distributional value | 0.0097 | 0.013 | 0.019 |
| ensemble NNs (4 ensembles) | peak value | 0.35 | 0.20 | 0.15 |
| | distributional value | 0.0049 | 0.0077 | 0.014 |

**Table S5. Parameters for the SDE model.**

To generate the training/test datasets, the values of 24 parameters were randomly picked from prespecified ranges (bold) and other parameters were fixed.

| Constant | Value | Description and source |
|---|---|---|
| $k_{MC}$ | **0.2-5 μM/h** | **MYC synthesis rate** |
| $k_S$ | 0.05 μM/h | EFm synthesis rate (serum) (Arbitrary value adjusted to match experimental observations presented here and [39,40]) |
| $k_{EFm}$ | **0.08-2.0 μM/h** | **EFm synthesis rate** |
| $k_b$ | **0.03-0.75 μM/h** | **EFp-independent EFm synthesis rate (serum)** |
| $k_{EFp}$ | **0.08-2.0 /h** | **E2F translation rate** |
| $k_{CD}$ | **0.01-0.1 μM/h** | **CYCD synthesis rate (MYC)** |
| $k_{CDS}$ | **0.1-2.0 μM/h** | **CYCD synthesis rate (serum)** |
| $k_{CE}$ | **0.07-1.5 μM/h** | **CYCE synthesis rate** |
| $k_{RB}$ | **0.05-0.9 μM/h** | **RB synthesis rate** |
| $k_{RE}$ | **36-360 /(μM*h)** | **RB-E2F formation rate** |
| $* k_{DP}$ | 3.6 μM/h | RB dephosphorylation rate[41] |
| $* k_{p1}$ | 18 /h | RB phosphorylation rate mediated by CYCD[41] |
| $* k_{p2}$ | 18 /h | RB phosphorylation rate mediated by CYCE[41] |
| $k_{AFb}$ | 0.007 μM/h | Basal ARF synthesis rate (Arbitrary value - included based on role in nucleolar integrity in absence of oncogenic stress) [42] |
| $k_{AFEF}$ | **0.003-0.075 μM/h** | **Synthesis rate of ARF by EFp** |
| $k_{AFMC}$ | **0.002-0.05 μM/h** | **Synthesis rate of ARF by MYC** |
| $k_{MREF}$ | **0.16-4.0 μM/h** | **Synthesis rate of miRNA by EFp** |
| $k_{MRMC}$ | **0.04-1.0 μM/h** | **Synthesis rate of miRNA by MYC** |
| $K_{AFMC}$ | **0.2-5.0 μM** | **Half-maximal MYC concentration (ARF synthesis)** |
| $K_{AFEF}$ | **0.1-2.5 μM** | **Half-maximal EFp concentration (ARF synthesis)** |
| $K_{MRMC}$ | **0.05-1.25 μM** | **Half-maximal MYC concentration (miRNA synthesis)** |
| $K_{MREF}$ | **0.05-1.25 μM** | **Half-maximal EFp concentration (miRNA synthesis)** |
| $K_{MC}$ | **0.03-0.75 μM** | **Half-maximal MYC concentration (EFm autoregulation)** |

| | | |
|---|---|---|
| $K_{MC1}$ | **0.5-12.5 μM** | **Half-maximal MYC concentration (EFp-independent EFm regulation)** |
| $K_S$ | **0.1%-2.5%** | **Half-maximal serum concentration** |
| $K_{EF}$ | **0.03-0.75 μM** | **Half-maximal EFp concentration (E2F autoregulation)** |
| $K_R$ | **20-200 μM** | **Half-maximal repression of EFm by MYC (Adjusted to match observations from this study)** |
| $K_{MR}$ | 0.6 μM | Half-maximal miRNA concentration (EFp repression) (Adjusted according to experimental observations [43]) |
| $K_{AFR}$ | **0.002-0.05 /μM** | **A constant to account for ARF-mediated EFp decay [44]** |
| $K_{RP}$ | **0.002-0.05 μM** | **Michaelis-Menten constant for constitutive dephosphorylation[41]** |
| $K_{CD}$ | 0.92 μM | Half-maximal CYCD concentration (RB phosphorylation)[45,46] |
| $K_{CE}$ | 0.92 μM | Half-maximal CYCE concentration (RB phosphorylation)[45,46] |
| $K_{MCCD}$ | 0.15 μM | Half-maximal MYC concentration (CYCD synthesis)[47] |
| $d_{EFm}$ | 0.25 /h | EFm decay constant [48,49] |
| $d_{EFp}$ | 0.35 /h | EFp decay constant [50] |
| $d_{CD}$ | 1.5 /h | CYCD decay constant[51,52] |
| $d_{CE}$ | 1.5 /h | CYCE decay constant[53,54] |
| $d_{RB}$ | 0.06 /h | RB decay constant[55] |
| $d_{RP}$ | 0.06 /h | Phospho-RB decay constant[55] (assume to be the same as $d_{RB}$) |
| $d_{RE}$ | 0.03 /h | RB-E2F decay constant[56] |
| $d_{MC}$ | 0.7 /h | MYC decay constant[57-59] |
| $d_{AF}$ | 0.12 /h | ARF decay constant [60-63] |
| $d_{MR}$ | 2.8 /h | miR-17-92 cluster miRNA decay constant[64] |

#### V.Supplemental figure legends

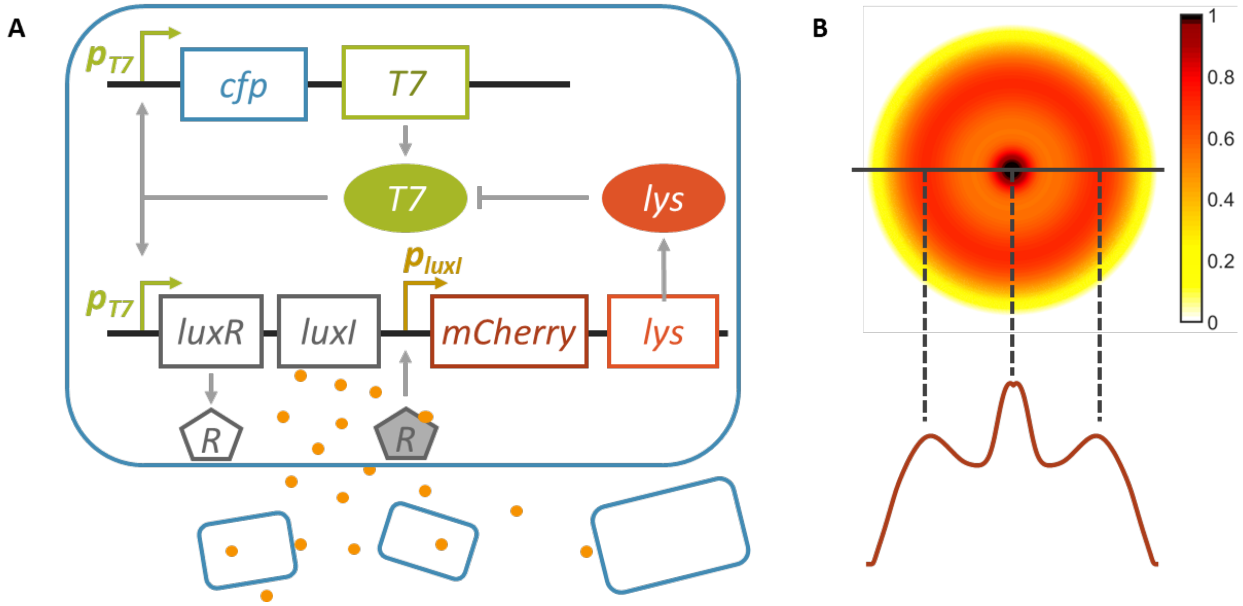


**Figure S1: PDE model description**

**A. A pattern-formation circuit.** The circuit consists of a T7 RNA polymerase that activates its own expression as well as the expression of LuxR and LuxI. Upon activation by T7RNAP (T7), LuxI mediates synthesis of AHL (orange dots), which can diffuse across the cell membrane. When the global AHL concentration surpasses a threshold, intracellular AHL binds to LuxR (R) to activate the synthesis of T7 lysozyme (lys). Lysozyme then binds to the T7RNAP and forms a T7-lysozyme complex, therefore inhibiting the T7RNAP binding to the T7 promoter. This complex also inhibits T7RNAP transcription.

**B. A schematic plot showing that the 1D concentration along a radius line is sufficient to represent the spherical geometry of the 2D pattern.** We choose hot colormap in Matlab and normalized the maximum concentration to be 1 for the plot. The 1D curves are used as ground truth of our model.

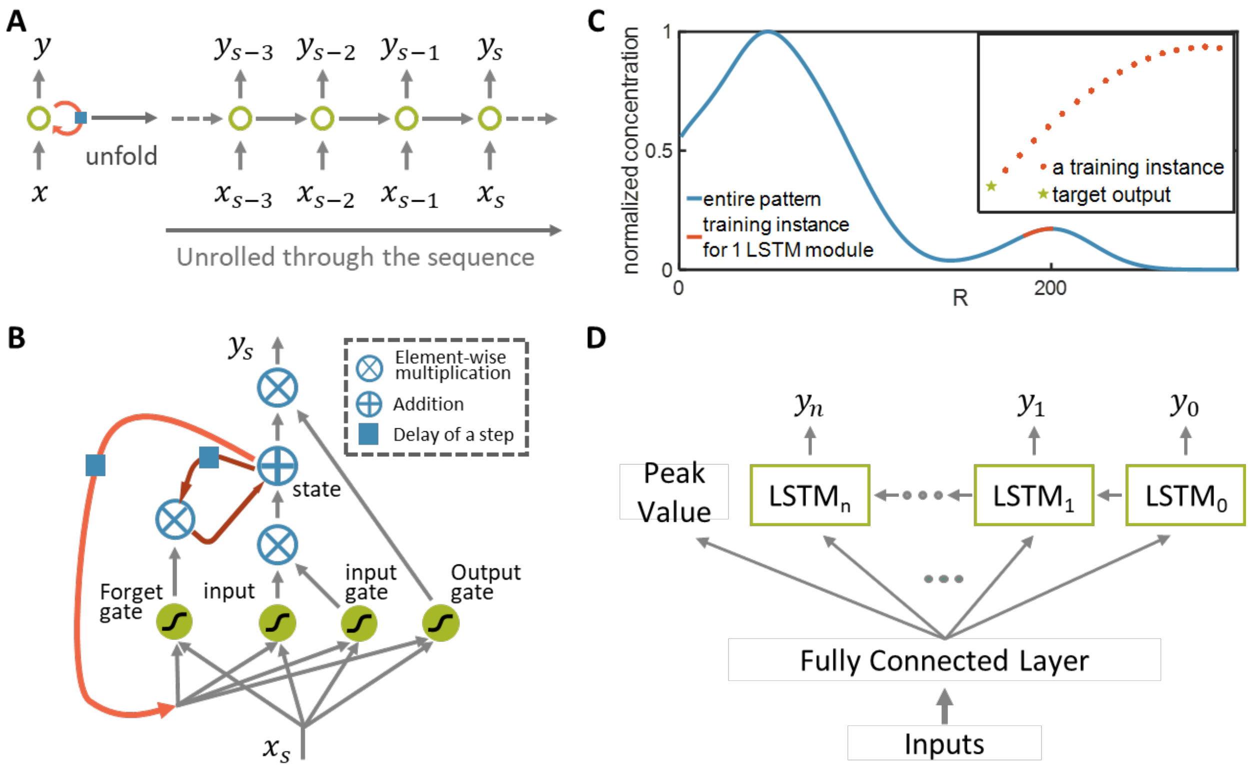


**Figure S2: Introduction to the concept of deep LSTM networks and the structure of the employed deep LSTM network.**

**A.** **A recurrent neuron.** A recurrent neuron receives a sequential input x, produces an output and sends that output back to itself. At each sequential step s (also called a frame), this recurrent neuron receives input $x_s$ as well as its own output from previous sequential step $y_{s-1}$. The black block indicates a delay of a single sequential step. This neuron (left) is the same as the unrolling computational graph (right), where each node is now associated with one particular sequential instance.

**B.** **A typical LSTM neural network unit.** In TensorFlow, LSTM cells can be simply implemented by using tf.contrib.rnn.BasicLSTMCell built-in function without needing to know the cell structure. In short, LSTM cells manage two state vectors, one is responsible for short-term memory and one is responsible for long-term memory. For each step, it adds some memories to long-term memories (controlled by input gate), drop some memories (controlled by the forget gate) and decide which parts of the long-term memories should be read and output at this step (controlled by the output gate). More details can be found at the referenced books[29,65].

**C.** **A training instance. We** discretize the x-axis into 501 points, so the entire pattern becomes a continuous pattern distribution series and is what we want to predict (blue line). Each point is associated with an LSTM module. There are 501 LSTM modules in total. For each module, the target output is a single value (green star), and the inputs are outputs of the previous *m* (*m*=16 in the figure demonstration, red dots) neighboring LSTM modules. Red line and the small figure window represent a training instance from that series for one LSTM module.

D. **The structure of the employed deep LSTM network.** The employed deep LSTM network consists of an input layer with inputs to be the parameters of mechanism-based model, a fully connected layer, LSTM arrays, and 2 output layers, one for predicting peak value of the distribution, one for predicting the normalized distribution.

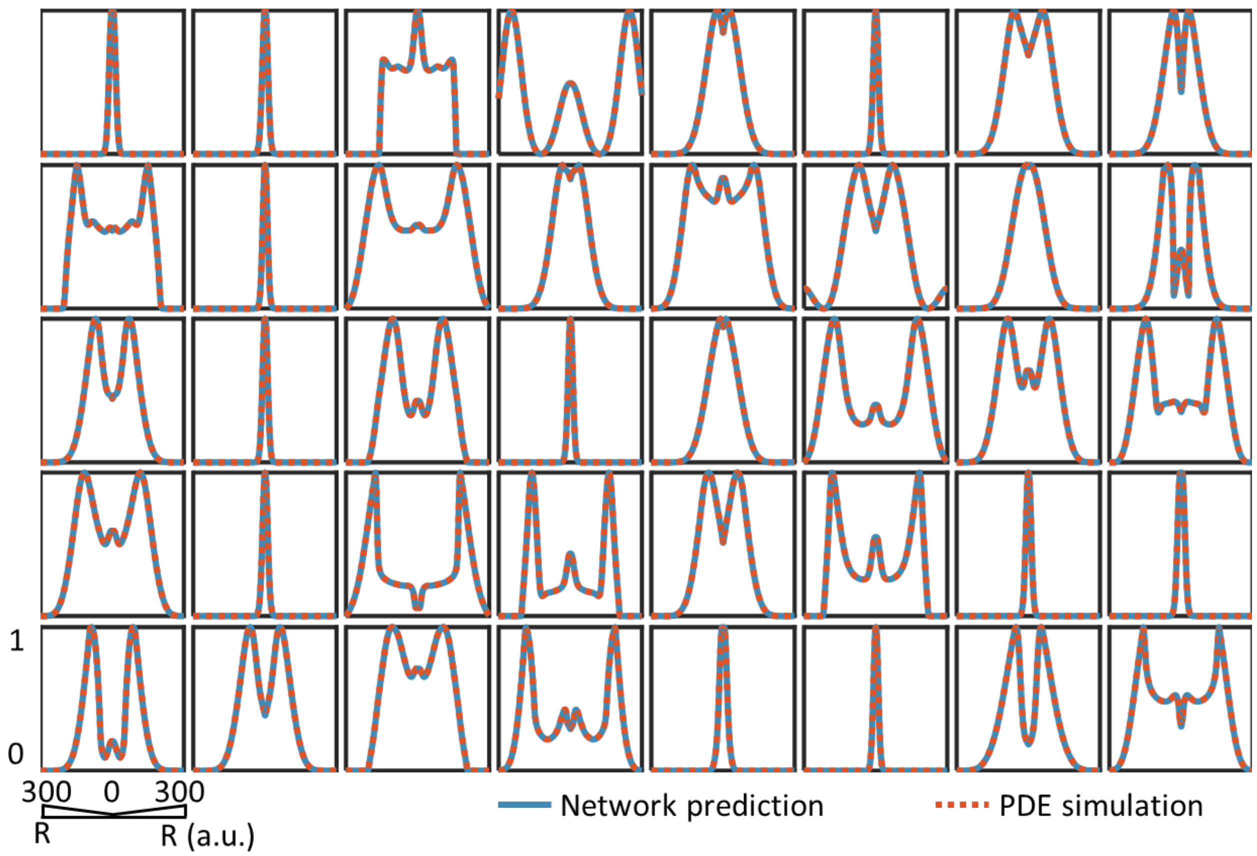


**Figure S3. Comparison between predicted distributions generated by neural network and distributions generated by mechanism-based model.** These examples are randomly selected from the training dataset.

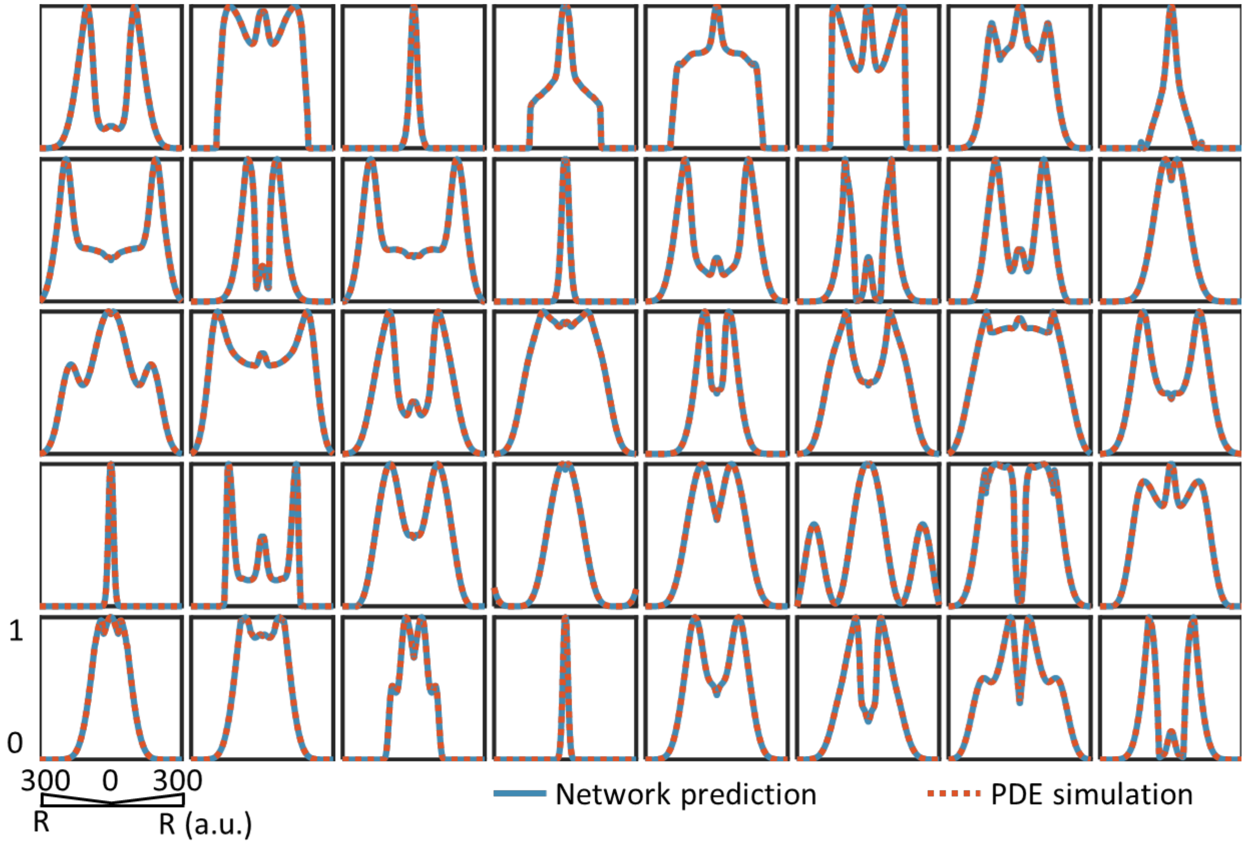


**Figure S4. Comparison between predicted distribution generated by neural network and distribution generated by mechanism-based model.** These are randomly selected from the test dataset, i.e., the dataset generated by mechanism-based model, however, never been used to train the neural networks.

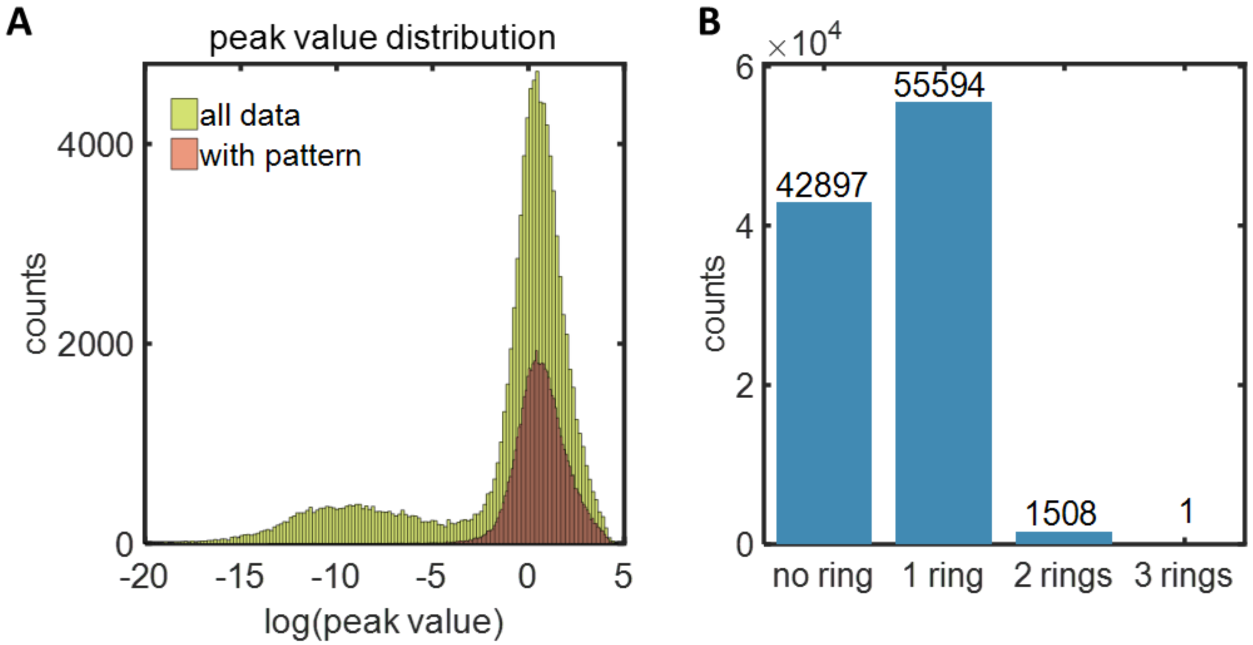


**Figure S5: Analysis of the training data acquired from simulation.**

**A. Peak value distribution.** The total training data size is $10^5$. The peak value for all the training data can be as low as $10^{-20}$, or as high as $10^5$. The peak value for data with pattern (one or more than one ring) is more concentrated at the higher end.

**B. Data pattern structure.** The training dataset consists of 42897 sets of data with no ring, 55594 sets of data with 1 ring, 1508 sets of data with 2 rings and only 1 set of data with 3 rings.

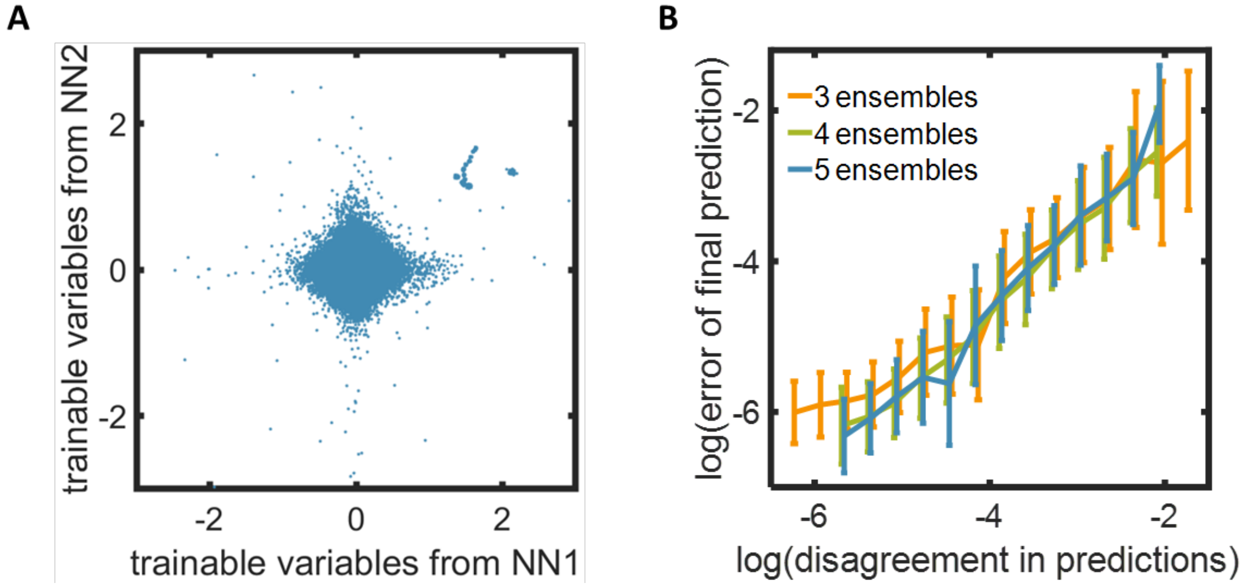


**Figure S6 Ensemble prediction analysis**

A. **Comparison of trainable variables (weights, bias) between 2 trained neural networks.** The difference in parameterization is due to random initialization and the properties of backpropagation.

B. **The increased disagreement in prediction is positively correlated with the increased error in predictions.** We tested the positive correlation between logarithm value of disagreement in prediction and logarithm value of error of final prediction using 3, 4, 5 ensembles of LSTM networks.

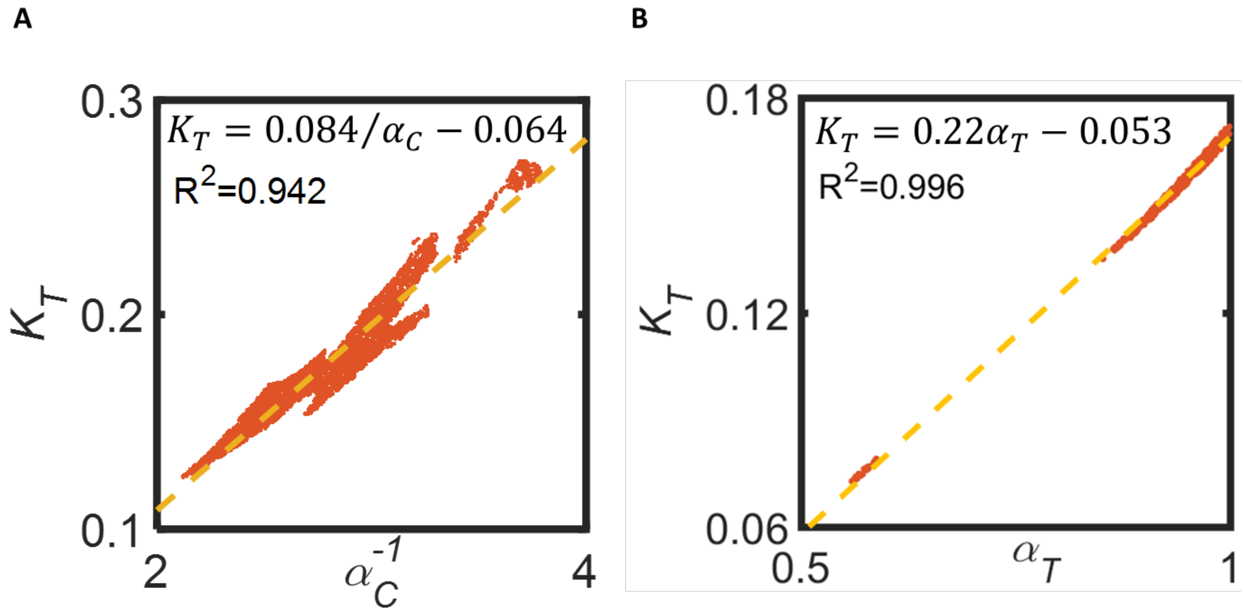


**Figure S7 Relationship of parameters to generate 3-ring patterns.** We screened through $10^8$ combinations of parameter sets using the ensemble prediction method, where we discarded predictions with disagreement in predictions larger than 0.1. For each of the screening, we vary two parameters of interest and fixed the rest and we only plot the parameter combination that can generate 3-rings. These NN predictions reveal the general criterion for generating 3-ring patterns.

**A. Negative relationship between cell growth rate on agar ($\boldsymbol{\alpha_C}$) and half activation constant of T7RNAP ($\boldsymbol{K_T}$).** If approximating that they are inversely proportional, we can get the fitting with $R^2 = 0.94$. Other parameters are fixed with constant values: $\alpha_A = 0.5$, $\alpha = 0.5$, $\beta = 0.5$, $K_\emptyset = 0.3$, $n = 0.5$, $\alpha_T = 0.8$, $\alpha_L = 0.3$, $K_C = 0.5$, $K_P = 0.5$, $d_A = 0.5$, $D = 1.0$.

**B. Linear correlation between half activation constant of T7RNAP ($\boldsymbol{K_T}$), and synthesis rate of T7RNAP ($\boldsymbol{\alpha_T}$) in order to generate 3-ring patterns.** $R^2 = 0.996$. Other parameters are fixed with constant values: $\alpha_A = 0.5$, $\alpha = 0.5$, $\beta = 0.5$, $K_\emptyset = 0.3$, $n = 0.5$, $\alpha_C = 0.5$, $\alpha_L = 0.3$, $K_C = 0.5$, $K_P = 0.5$, $d_A = 0.5$, $D = 1.0$.

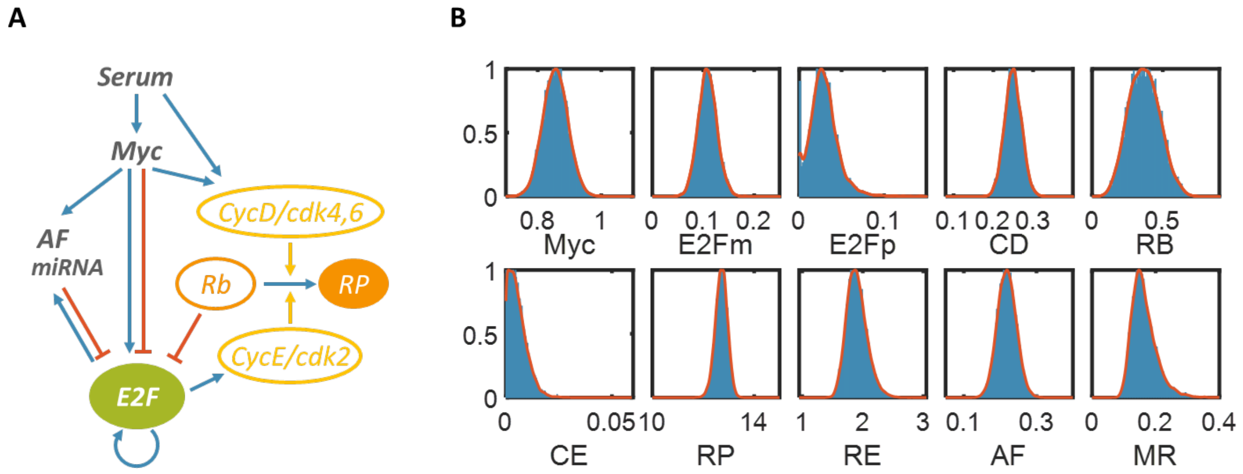


**Figure S8 Stochastic Myc-E2F pathway in cell-cycle progression.** This model is from Jeffrey Wong, et al[66].

**A. Diagram of a concrete mechanism-based SDE model example.** E2F functions as the output of the Rb-E2F signaling pathway and is involved in multiple positive-feedback loops (Fig. 1a). In quiescent cells, E2F is bound to and repressed by Rb. With sufficient growth stimulation, phosphorylation by Myc-induced *cyclin D* (CycD) - Cdk4,6 removes Rb repression; Myc also induces E2F transcription. Subsequently, E2F activates the transcription of CycE, which forms a complex with Cdk2 to further remove Rb repression by phosphorylation, establishing a positive-feedback loop. E2F also activates its own transcription, constituting another positive-feedback loop.

**B. Histogram of stochastic simulations.** $10^4$ stochastic simulations are used to make this plot. We split the data into 100 bins and plot the histogram with no gap between bars. With a sufficiently large number of simulations, this distribution converges to an approximately continuous curve. The red dotted curve is the kernel fitting using Matlab function fitdist() function, which will be used as the ground truth to train the neural network.

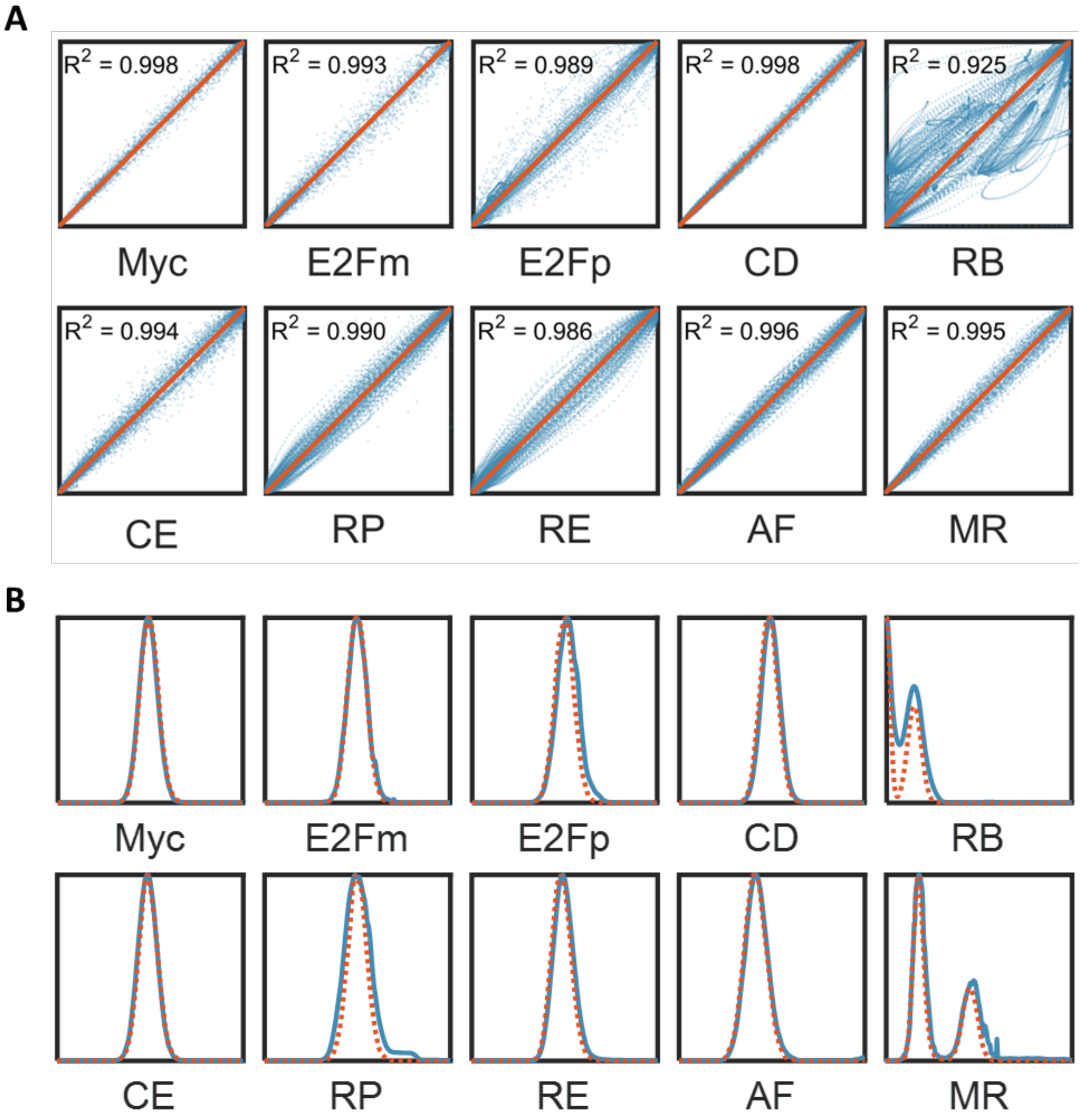


**Figure S9. Neural network performance and sample prediction.**

A. **Accuracy analysis for deep LSTM network prediction system.** On the top is the plot with the predicted distribution by neural network against the distribution generated by numerical simulation. On the bottom is the plot with the peak value generated by neural networks against the peak value generated by numerical simulations. Perfect alignment corresponds to the $y = x$ line. We calculated the R-square to measure how close they are. The test sample size is s (=10,000). For each of the distribution, there are 1,000 discrete points representing space segregation.

**B. Representative distributional sample predicted by neural network.** Blue lines are the predicted distributions generated by trained neural network, red dashed lines are the distributions generated by numerical simulations.

#### VI.Supplemental References


1 Payne, S. *et al.* Temporal control of self-organized pattern formation without morphogen gradients in bacteria. *Mol Syst Biol* **9**, 697, doi:10.1038/msb.2013.55 (2013).

2 Kenkre, V. M. Results from variants of the Fisher equation in the study of epidemics and bacteria. *Physica A* **342**, 242-248, doi:DOI 10.1016/j.physa.2004.04.084 (2004).

3 Murray, J. D., Cook, J., Tyson, R. & Lubkin, S. R. Spatial pattern formation in biology: I. Dermal wound healing. II. Bacterial patterns. *J Franklin I* **335B**, 303-332, doi:Doi 10.1016/S0016-0032(97)00034-3 (1998).

4 Maini, P. K. Using mathematical models to help understand biological pattern formation. *Comptes rendus biologies* **327**, 225-234 (2004).

5 Kumar, A. & Patel, S. S. Inhibition of T7 RNA polymerase: Transcription initiation and transition from initiation to elongation are inhibited by T7 lysozyme via a ternary complex with RNA polymerase and promoter DNA. *Biochemistry-Us* **36**, 13954-13962, doi:Doi 10.1021/Bi971432y (1997).

6 Yao, G., Lee, T. J., Mori, S., Nevins, J. R. & You, L. C. A bistable Rb-E2F switch underlies the restriction point. *Nat Cell Biol* **10**, 476-U255, doi:10.1038/ncb1711 (2008).

7 Lee, T. J., Yao, G. A., Bennett, D. C., Nevins, J. R. & You, L. C. Stochastic E2F Activation and Reconciliation of Phenomenological Cell-Cycle Models. *Plos Biol* **8**, doi:10.1371/journal.pbio.1000488 (2010).

8 Gillespie, D. T. The chemical Langevin equation. *J Chem Phys* **113**, 297-306, doi:Doi 10.1063/1.481811 (2000).

9 Forsyth, D. & Ponce, J. *Computer vision : a modern approach.* (Prentice Hall, 2003).

10 He, K., Zhang, X., Ren, S. & Sun, J. Spatial Pyramid Pooling in Deep Convolutional Networks for Visual Recognition. *IEEE Trans Pattern Anal Mach Intell* **37**, 1904-1916, doi:10.1109/TPAMI.2015.2389824 (2015).

11 Goodfellow, I., Bengio, Y. & Courville, A. Deep Generative Models. *Adapt Comput Mach Le*, 645-710 (2016).

12 Jaitly, N. & Hinton, G. Learning a Better Representation of Speech Soundwaves Using Restricted Boltzmann Machines. *2011 Ieee International Conference on Acoustics, Speech, and Signal Processing*, 5884-5887 (2011).

13 Graves, A., Mohamed, A. R. & Hinton, G. Speech Recognition with Deep Recurrent Neural Networks. *2013 Ieee International Conference on Acoustics, Speech and Signal Processing (Icassp)*, 6645-6649 (2013).

14 Sainath, T. N., Mohamed, A. R., Kingsbury, B. & Ramabhadran, B. Deep Convolutional Neural Networks for Lvcsr. *2013 Ieee International Conference on Acoustics, Speech and Signal Processing (Icassp)*, 8614-8618 (2013).

15 Lu, L., Zhang, X. X., Cho, K. H. & Renals, S. A Study of the Recurrent Neural Network Encoder-Decoder for Large Vocabulary Speech Recognition. *16th Annual Conference of the International Speech Communication Association (Interspeech 2015), Vols 1-5*, 3249-3253 (2015).

16 Gibney, E. Google AI algorithm masters ancient game of Go. *Nature* **529**, 445-446, doi:10.1038/529445a (2016).

17 Silver, D. *et al.* Mastering the game of Go with deep neural networks and tree search. *Nature* **529**, 484-489, doi:10.1038/nature16961 (2016).

18 Silver, D. *et al.* Mastering the game of Go without human knowledge. *Nature* **550**, 354-359, doi:10.1038/nature24270 (2017).

19 Alipanahi, B., Delong, A., Weirauch, M. T. & Frey, B. J. Predicting the sequence specificities of DNA- and RNA-binding proteins by deep learning. *Nat Biotechnol* **33**, 831-838, doi:10.1038/nbt.3300 (2015).

20 LeCun, Y., Bengio, Y. & Hinton, G. Deep learning. *Nature* **521**, 436-444, doi:10.1038/nature14539 (2015).

21 Putin, E. *et al.* Deep biomarkers of human aging: Application of deep neural networks to biomarker development. *Aging (Albany NY)* **8**, 1021-1033 (2016).

22 Ananthakrishnan, A. N. *et al.* Gut Microbiome Function Predicts Response to Anti-integrin Biologic Therapy in Inflammatory Bowel Diseases. *Cell Host Microbe* **21**, 603-610 e603, doi:10.1016/j.chom.2017.04.010 (2017).

23 Esteva, A. *et al.* Dermatologist-level classification of skin cancer with deep neural networks. *Nature* **542**, 115-118, doi:10.1038/nature21056 (2017).

24 Altae-Tran, H., Ramsundar, B., Pappu, A. S. & Pande, V. Low Data Drug Discovery with One-Shot Learning. *ACS Cent Sci* **3**, 283-293 (2017).

25 Camacho, D. M., Collins, K. M., Powers, R. K., Costello, J. C. & Collins, J. J. Next-Generation Machine Learning for Biological Networks. *Cell* **173**, 1581-1592, doi:10.1016/j.cell.2018.05.015 (2018).

26 Ryu, J. Y., Kim, H. U. & Lee, S. Y. Deep learning improves prediction of drug-drug and drug-food interactions. *P Natl Acad Sci USA* **115**, E4304-E4311 (2018).

27 Rumelhart, D. E., Hinton, G. E. & Williams, R. J. Learning Representations by Back-Propagating Errors. *Nature* **323**, 533-536, doi:DOI 10.1038/323533a0 (1986).

28 Géron, A. l. *Hands-on machine learning with Scikit-Learn and TensorFlow : concepts, tools, and techniques to build intelligent systems*. First edition. edn, (O'Reilly Media, Inc., 2017).

29 Goodfellow, I., Bengio, Y. & Courville, A. Deep Learning. *Adapt Comput Mach Le*, 1-775 (2016).

30 Pascanu, R., Mikolov, T. & Bengio, Y. in *Proceedings of the 30th International Conference on Machine Learning* Vol. 28 (eds Dasgupta Sanjoy & McAllester David) 1310--1318 (PMLR, Proceedings of Machine Learning Research, 2013).

31 Hochreiter, S. & Schmidhuber, J. Long short-term memory. *Neural Comput* **9**, 1735-1780 (1997).

32 Graves, A. & Schmidhuber, J. Framewise phoneme classification with bidirectional LSTM and other neural network architectures. *Neural Netw* **18**, 602-610, doi:10.1016/j.neunet.2005.06.042 (2005).

33 Vinyals, W. Z. a. I. S. a. O. Recurrent Neural Network Regularization. (2015).

34 He, K. M., Zhang, X. Y., Ren, S. Q. & Sun, J. Delving Deep into Rectifiers: Surpassing Human-Level Performance on ImageNet Classification. *Ieee I Conf Comp Vis*, 1026-1034, doi:10.1109/Iccv.2015.123 (2015).

35 Song, H., Payne, S., Gray, M. & You, L. Spatiotemporal modulation of biodiversity in a synthetic chemical-mediated ecosystem. *Nat Chem Biol* **5**, 929-935, doi:10.1038/nchembio.244 (2009).

36 You, L., Cox, R. S., 3rd, Weiss, R. & Arnold, F. H. Programmed population control by cell-cell communication and regulated killing. *Nature* **428**, 868-871, doi:10.1038/nature02491 (2004).

37 Collins, C. H., Leadbetter, J. R. & Arnold, F. H. Dual selection enhances the signaling specificity of a variant of the quorum-sensing transcriptional activator LuxR. *Nature biotechnology* **24**, 708-712, doi:10.1038/nbt1209 (2006).

38 Aronson, D., Weinberger, H. & Multidimensional nonlinear diffusion arising in population genetics. *Advances in Mathematics* **30.1**, 33-76 (1978).

39 Hsiao, K. M., McMahon, S. L. & Farnham, P. J. Multiple DNA elements are required for the growth regulation of the mouse E2F1 promoter. *Genes Dev* **8**, 1526-1537 (1994).

40 Smith, E. J., Leone, G., DeGregori, J., Jakoi, L. & Nevins, J. R. The accumulation of an E2F-p130 transcriptional repressor distinguishes a G0 cell state from a G1 cell state. *Mol Cell Biol* **16**, 6965-6976 (1996).

41 Kholodenko, B. N. Cell-signalling dynamics in time and space. *Nat Rev Mol Cell Biol* **7**, 165-176, doi:10.1038/nrm1838 (2006).

42 Apicelli, A. J. *et al.* A non-tumor suppressor role for basal p19ARF in maintaining nucleolar structure and function. *Mol Cell Biol* **28**, 1068--1080 (2008).

43 Pickering, M. T., Stadler, B. M. & Kowalik, T. F. miR-17 and miR-20a temper an E2F1-induced G1 checkpoint to regulate cell cycle progression. *Oncogene* (2008).

44 Martelli, F. *et al.* p19ARF targets certain E2F species for degradation. *Proc Natl Acad Sci U S A* **98**, 4455-4460, doi:10.1073/pnas.081061398 (2001).

45 Pan, W., Sun, T., Hoess, R. & Grafstrom, R. Defining the minimal portion of the retinoblastoma protein that serves as an efficient substrate for cdk4 kinase/cyclin D1 complex. *Carcinogenesis* **19**, 765-769 (1998).

46 Grafstrom, R. H., Pan, W. & Hoess, R. H. Defining the substrate specificity of cdk4 kinase-cyclin D1 complex. *Carcinogenesis* **20**, 193-198 (1999).

47 Park, S. *et al.* Determination of binding constant of transcription factor myc-max/max-max and E-box DNA: the effect of inhibitors on the binding. *Biochim Biophys Acta* **1670**, 217-228, doi:10.1016/j.bbagen.2003.12.007 (2004).

48 Bartley, S. M., Szakaly, R. J. & Farnham, P. J. Characterization of the 3' untranslated region of mouse E2F1 mRNA. *Gene* **223**, 355-360 (1998).

49 Saunders, N. A., Dicker, A. J., Jones, S. J. & Dahler, A. L. E2F1 messenger RNA is destabilized in response to a growth inhibitor in normal human keratinocytes but not in a squamous carcinoma cell line. *Cancer Res* **58**, 1646-1649 (1998).

50 Helin, K. Regulation of cell proliferation by the E2F transcription factors. *Curr Opin Genet Dev* **8**, 28-35, doi:S0959-437X(98)80058-0 [pii] (1998).

51 Diehl, J. A., Zindy, F. & Sherr, C. J. Inhibition of cyclin D1 phosphorylation on threonine-286 prevents its rapid degradation via the ubiquitin-proteasome pathway. *Genes Dev* **11**, 957-972 (1997).

52 Sherr, C. J. & Roberts, J. M. CDK inhibitors: positive and negative regulators of G1-phase progression. *Genes Dev* **13**, 1501-1512 (1999).

53 Clurman, B. E., Sheaff, R. J., Thress, K., Groudine, M. & Roberts, J. M. Turnover of cyclin E by the ubiquitin-proteasome pathway is regulated by cdk2 binding and cyclin phosphorylation. *Genes Dev* **10**, 1979-1990 (1996).

54 Won, K. A. & Reed, S. I. Activation of cyclin E/CDK2 is coupled to site-specific autophosphorylation and ubiquitin-dependent degradation of cyclin E. *EMBO J* **15**, 4182-4193 (1996).

55 Mihara, K. *et al.* Cell cycle-dependent regulation of phosphorylation of the human retinoblastoma gene product. *Science* **246**, 1300-1303 (1989).

56 Buchler, N. E., Gerland, U. & Hwa, T. Nonlinear protein degradation and the function of genetic circuits. *Proc Natl Acad Sci U S A* **102**, 9559-9564, doi:10.1073/pnas.0409553102 (2005).

57 Sears, R., Leone, G., DeGregori, J. & Nevins, J. R. Ras enhances Myc protein stability. *Mol Cell* **3**, 169-179 (1999).

58 Sears, R. *et al.* Multiple Ras-dependent phosphorylation pathways regulate Myc protein stability. *Genes Dev* **14**, 2501-2514 (2000).

59 Yeh, E. *et al.* A signalling pathway controlling c-Myc degradation that impacts oncogenic transformation of human cells. *Nat Cell Biol* **6**, 308-318, doi:10.1038/ncb1110 (2004).

60 Bertwistle, D., Sugimoto, M. & Sherr, C. J. Physical and functional interactions of the Arf tumor suppressor protein with nucleophosmin/B23. *Mol Cell Biol* **24**, 985-996 (2004).

61 Colombo, E. *et al.* Nucleophosmin is required for DNA integrity and p19Arf protein stability. *Mol Cell Biol* **25**, 8874-8886, doi:10.1128/MCB.25.20.8874-8886.2005 (2005).

62 Kuo, M. L., den Besten, W., Bertwistle, D., Roussel, M. F. & Sherr, C. J. N-terminal polyubiquitination and degradation of the Arf tumor suppressor. *Genes Dev* **18**, 1862-1874, doi:10.1101/gad.1213904 (2004).

63 Pollice, A. *et al.* Functional and physical interaction of the human ARF tumor suppressor with Tat-binding protein-1. *J Biol Chem* **279**, 6345-6353, doi:10.1074/jbc.M310957200 (2004).

64 O'Donnell, K. A., Wentzel, E. A., Zeller, K. I., Dang, C. V. & Mendell, J. T. c-Myc-regulated microRNAs modulate E2F1 expression. *Nature* **435**, 839--843 (2005).

65 Géron, A. l. *Hands-on machine learning with Scikit-Learn and TensorFlow : concepts, tools, and techniques to build intelligent systems*. First edition. edn.

66 Wong, J. V., Yao, G., Nevins, J. R. & You, L. Viral-mediated noisy gene expression reveals biphasic E2f1 response to MYC. *Mol Cell* **41**, 275-285, doi:10.1016/j.molcel.2011.01.014 (2011).
